# Mimics of the chordate gut–brain hormone neurotensin in parasitic intestinal hookworms

**DOI:** 10.64898/2026.01.24.701534

**Authors:** Thomas Lund Koch, Ho Yan Yeung, Iris Bea L. Ramiro, Joshua P. Torres, Alina Anhnhi Vo, Seph Palomino, Ebbe Engholm, Samuel D. Robinson, Lachlan D. Rash, Zildjian G. Acyatan, Nicholas C. Schumann, Kasper B. Pedersen, Knud J. Jensen, Katrine T. Schjoldager, Amol Patwardhan, Neil D. Young, Michael J. Robertson, Helena Safavi-Hemami

## Abstract

Hookworms of the family *Ancylostomatidae* are intestinal parasites that infect hundreds of millions of people worldwide, contributing to malnutrition, anemia, and impaired development. While hookworms are known to secrete immunomodulatory molecules to evade host defenses, it has been unclear whether they also exploit host hormonal signaling. Here, we identify a previously unrecognized family of hookworm peptides, which we term ancylotensins. Ancylotensins share strong sequence similarity with the chordate hormone neurotensin, a key regulator of metabolism, gut–brain communication, and intestinal function. *In vitro* pharmacological assays, *ex vivo* gut contractility studies, and cryo-electron microscopy demonstrate that ancylotensins closely replicate both the structural and functional properties of mammalian neurotensin to modulate gut function. Furthermore, ancylotensins do not share common ancestry with chordate neurotensin and instead evolved independently via convergent evolution. These findings reveal the existence of gut peptide mimicry as a mechanism by which intestinal parasites can manipulate host physiology.

## Introduction

Hookworms of the family Ancylostomatidae are nematode parasites that currently infect nearly 500 million people worldwide^1,2^. The primary human-infecting species are *Necator americanus* and *Ancylostoma duodenale*, although zoonotic species such as *Ancylostoma ceylanicum* and *Ancylostoma caninum* can also infect humans^1^. Infective third-stage larvae typically develop in soil and enter a new host either by penetrating the skin or through ingestion. After entry, they migrate through the lungs before establishing in the small intestine, where adult worms feed on blood, mate, and release eggs that are passed in feces (Figure 1A). Individual worms can persist in this niche for several years, which can cause iron-deficiency anemia and impair growth and cognition, particularly in children and women of reproductive age^1^.

**Figure 1.**
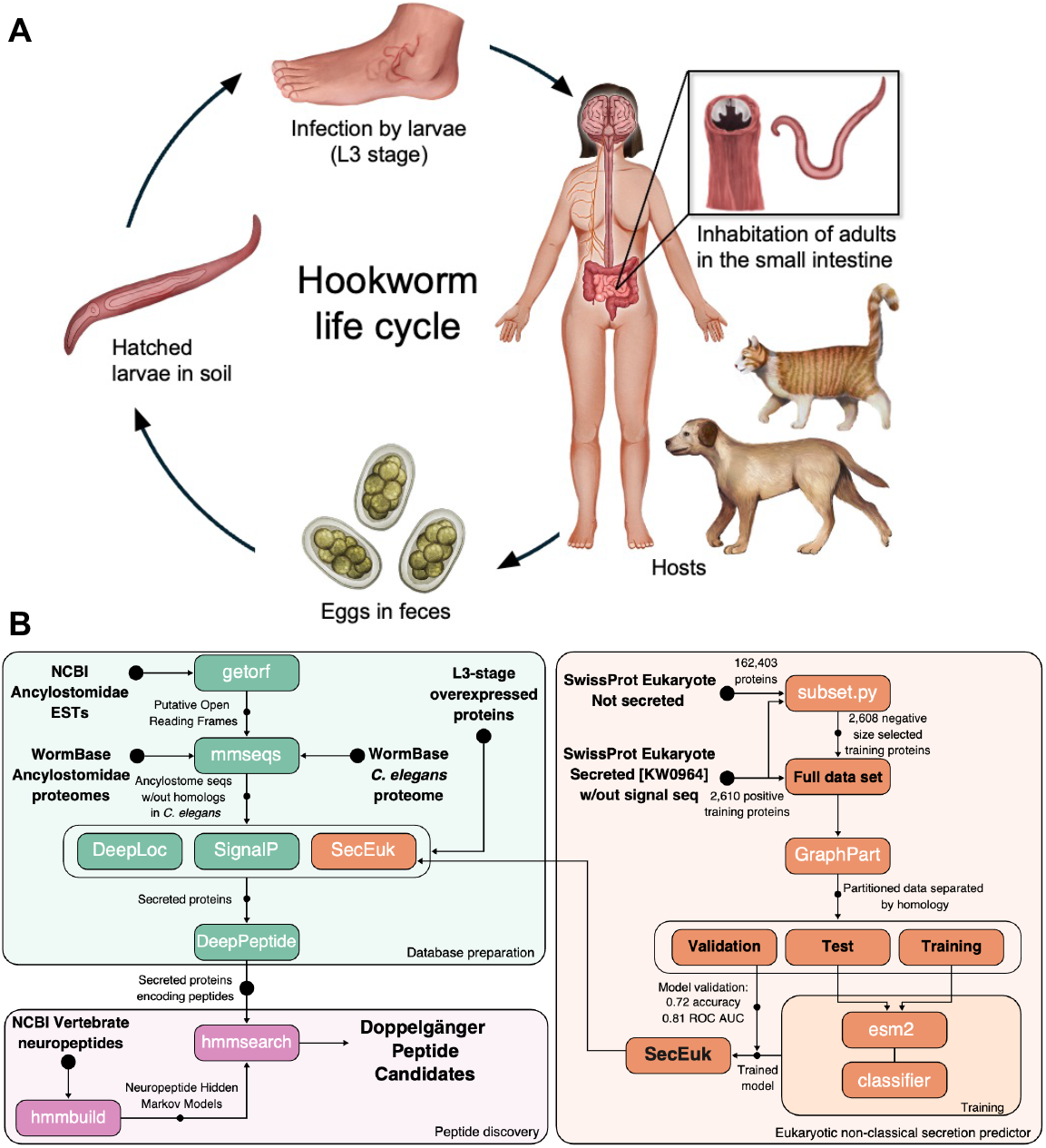
Intestinal hookworms of the family *Ancylostomatidae* express molecular mimics of host gut hormones. (A) Schematic of the hookworm life cycle. Illustration by ilusea studio. (B) Computational pipeline used for the discovery of doppelgänger peptide candidates from hookworm secretomes.

A key factor in hookworm survival is the secretion of parasite-derived proteins and peptides (the hookworm secretome) which includes anticoagulants^3^ and immunomodulators^4^ essential for blood feeding and immune evasion, respectively. However, many secreted molecules may remain undetected or uncharacterized, representing both a gap in understanding host-parasite interactions and a potential resource of bioactive compounds. Here, we hypothesized that hookworms, through their long-term adaptation to and residence in the host intestine, may have evolved peptides that mimic gut hormones to manipulate host physiology. We refer to such molecular mimics—peptides that resemble the peptide hormones of another organism—as doppelgänger peptides. We have previously shown that doppelgänger peptides are typically short and share only limited sequence similarity with their hormone counterparts, making them challenging to detect^5^. To overcome this, we generated hidden Markov models (HMMs) for known human neuropeptides and peptide hormones and systematically interrogated hookworm secretomes. This approach identified multiple candidates, including ancylotensins, secreted hookworm peptides that mimic the structure and activity of the host hormone neurotensin.

Neurotensin is a key vertebrate hormone involved in metabolic regulation^6^, intestinal function^7^ and gut-brain communication^8,9^. Secreted from intestinal N cells in response to dietary fat, neurotensin slows gastric emptying, promotes lipid absorption, and modulates gut motility and blood flow^10^. Beyond these local effects, neurotensin signals to the brain both endocrinologically, by crossing the blood-brain barrier, and neuronally, via binding sites on sensory and visceromotor components of the vagus nerve^8,11^. Neurotensin exerts its effects primarily through two class A G protein-coupled receptors (GPCRs), neurotensin receptor 1 (NTSR1) and neurotensin receptor 2 (NTSR2), and interacts with neurotensin receptor 3 (NTSR3, also known as sortilin), a member of the Vps10p-domain receptor family. A shorter peptide resembling the neurotensin C-terminus, neuromedin N, is encoded on the same precursor and co-secreted with neurotensin, sharing many of its receptor interactions^12,13^. Both peptides share a conserved Pro-Tyr-Ile-Leu (PYIL) motif important for receptor engagement^14^. Given its role in coordinating nutrient processing and potential role in central regulation of satiety and energy balance, neurotensin has been investigated as a therapeutic target for metabolic disorders^9,15^. In addition, neurotensin modulates central pain pathways, with several neurotensin analogs under investigation as analgesics^16,17^.

Here, using genomics, pharmacological, and structural analyses, we show that hookworms have evolved neurotensin-like peptides that can act on the neurotensin signaling system of their mammalian hosts. Our findings demonstrate the convergent evolution of host-like neuropeptides in gut-inhabiting nematodes, suggesting host hormone mimicry as a strategy in hookworm parasitism.

## Results

### Multiple doppelgänger peptide candidates are present in Ancylostomatidae

We compiled secreted-protein repertoires from four hookworm species (*N. americanus, A. duodenale, A. ceylanicum*, and *A. caninum*), which are classified as Clade V nematodes^18^, to search for doppelgänger peptides of mammalian neuropeptides. Predicted proteins (WormBase ParaSite and NCBI) and open reading frames below 400 amino acids derived from expressed sequence tags (ESTs) were compared to proteins from the free-living nematode *Caenorhabditis elegans* (also a Clade V nematode) to retain only proteins unique to *Ancylostomidae*. For *N. americanus* and *A. ceylanicum*, we further extracted secreted products overexpressed in the infectious L3 stage^19,20^. These proteins were filtered for classical and non-classical secretion with multiple tools^20-22^ to keep only products that are predicted to be secreted from the nematodes. We next used DeepPeptide^23^ to identify the subset that are predicted to encode mature peptides (Figure 1B, File S1-S2). This database was finally queried with HMMs trained on human neuropeptides and peptide hormones to identify hookworm secretome products with high similarity to human peptides. The HMM-identified candidates were then subjected to DeepPeptide^23^ precursor processing predictions together with manual inspection for bioactive peptide motifs and proteolytic cleavage sites, ultimately yielding two doppelgänger peptide candidates: a neurotensin/neuromedin N-like and a kisspeptin-like peptide from hookworms (Figure S1). Of these, the peptide showing similarity to neurotensin/neuromedin N was prioritized for subsequent analyses, as it most closely resembled the chordate hormones in both sequence and predicted processing and contained the conserved motif required for receptor engagement. Pharmacological data for the kisspeptin-like peptide obtained from the National Institute of Mental Health’s Psychoactive Drug Screening Program (PDSP) at the University of North Carolina at Chapel Hill is provided in Figure S2.

### Doppelgänger peptides of neurotensin are found in multiple hookworm species

Following identification of the neurotensin/neuromedin N-like doppelgänger candidate, we used this sequence as a query to search additional hookworm genome and transcriptome assemblies.

This analysis uncovered multiple similar sequences across other ancylostome species (Figure 2A, Figure S3, File S3), hereafter referred to as ancylotensins. All identified ancylotensin precursors encode peptides that share the C-terminal PYIL motif characteristic of neurotensin/neuromedin-N peptides with most also containing a positively charged residue immediately upstream of the PYIL motif. Moreover, like neurotensin, ancylotensins begin with an N-terminal Glu or Gln residue predicted to cyclize to pyroglutamate, a modification that confers resistance to proteolysis (Figure 2A). Unlike in vertebrates, where neurotensin and neuromedin N occur as a tandem pair on the same precursor (Figure 2B), hookworm ancylotensin genes encode multiple peptide repeats (Figure 2C). Each repeat is flanked by predicted basic (Arg, R) or dibasic (Lys-Arg, KR) cleavage sites (Figure 2A, Figure S3), a hallmark of neuropeptide processing. Given that neurotensin signaling systems are confined to chordates^24^ and absent from protostomes such as hookworms, we hypothesized that ancylotensins do not share common ancestry with the chordate peptide hormone but instead evolved convergently (Figure 2D) to hijack the neurotensin signaling system.

**Figure 2.**
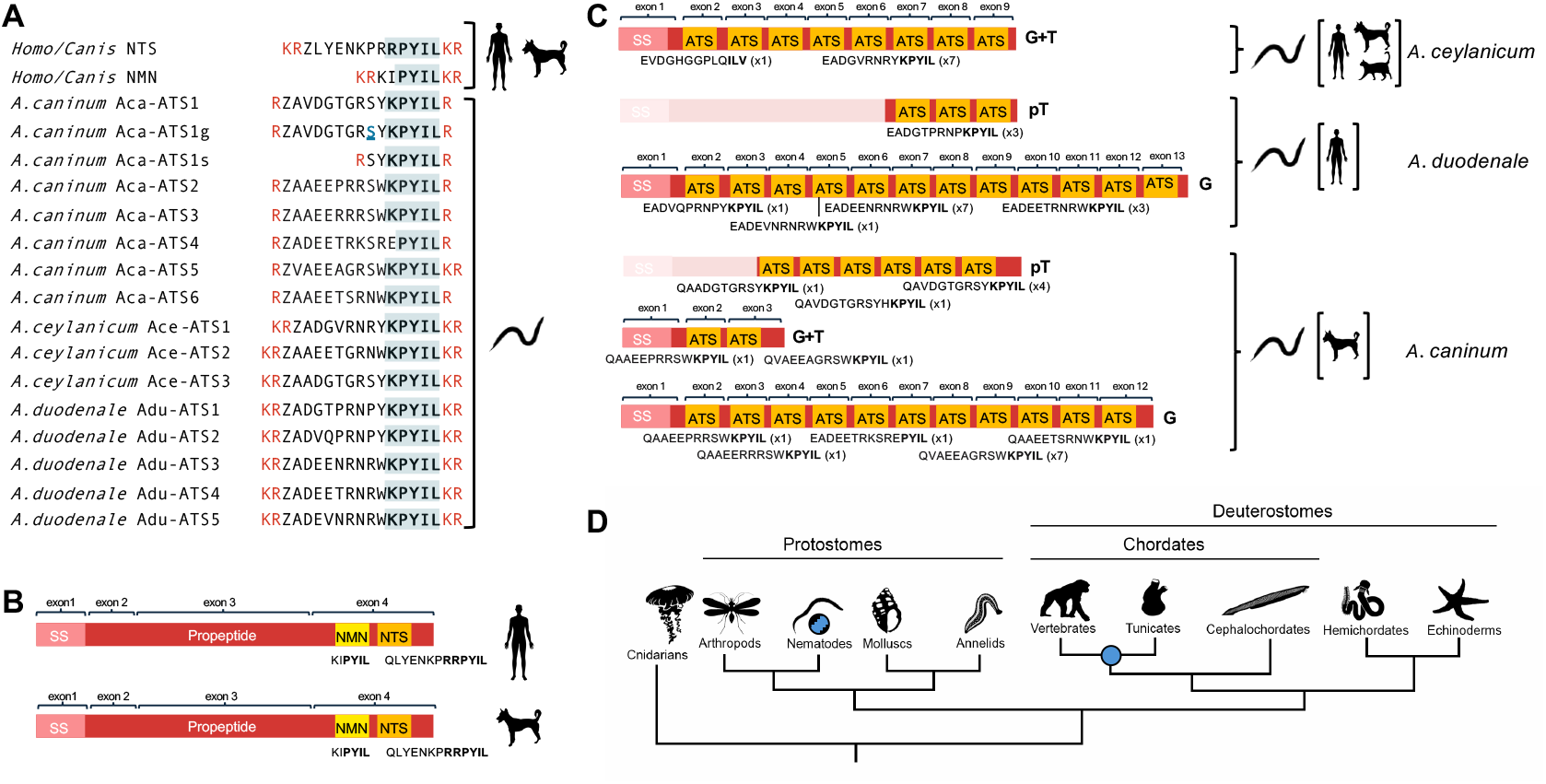
Ancylotensin (ATS) sequences identified from hookworm transcriptomes and genomes. (A) Predicted mature sequences of all ancylotensins identified in three hookworm species (*Ancylostoma caninum, Ancylostoma ceylanicum*, and *Ancylostoma duodenale*), aligned with human and dog neurotensin (NTS) and neuromedin (NMN). Flanking amino acids representing proteolytic cleavage sites are shown in red. Conserved residues are highlighted. Z=pyroglutamic acid, <u>S</u>= predicted *O*-glycosylation site. (B) Protein precursor organization of NTS/NMN in human and dog, and (C) of ATS in hookworms. Regions encoding the mature peptides are shown in yellow (NMN) and orange (NTS and ATS). Signal sequences are depicted in pink and propeptide regions in red. Exon positions are indicated above precursors, and peptide sequences and copy numbers are shown below. Shaded areas indicate incomplete sequence information. Letters on the right of protein precursors indicate if sequences derived from genes (G), transcripts (T), or partial transcripts (pT). (D) Cladogram showing that the NTS/NMN signaling system is confined to chordates (blue filled circle) and that NTS/NMN mimics (ancylotensins) in parasitic nematodes evolved independently via convergent evolution (blue striped circles).

### Evolutionary analysis reveals the convergent evolution of ancylotensins

Ancylotensin genes are present in the genomes of three hookworm species examined here (*A. duodenale, A. ceylanicum*, and *A. caninum)*, with transcription supported by RNA-seq and EST data (Table S1). By contrast, no ancylotensin homologs were detected in other closely related species, including *N. americanus* (family *Ancylostomidae*), suggesting that this is a recently evolved, lineage-specific gene family. In *A. duodenale* and *A. caninum*, ancylotensin-encoding genes reside on homologous chromosome IV (based *on A. duodenale* naming) and are syntenic between the two species (Figure S4A-B). By comparison, although the *A. ceylanicum* gene also maps to the homologous chromosome, its surrounding genomic neighborhood shows little synteny (Figure S4B). In *A. caninum* we identified two adjacent ancylotensin genes: one encoding the characteristic tandem peptide repeats found in other species, and one encoding only two ancylotensin peptides (Figure 2C). *A. duodenale* also carries a second ancylotensin gene, located on chromosome II, which appears to be unique to this species.

Ancylotensin genes are characterized by a highly repetitive architecture, typically consisting of 10–16 exons (Figure 2C, Figure S4C). With the exception of the first exon, each exon encodes a single ancylotensin peptide. These peptide-encoding exons are nearly identical within each species and show much higher similarity within than between species (Figure S5). This pattern is consistent with concerted evolution, a scenario in which repeated units (exons or whole genes) within a gene family are homogenized over time. Mechanisms such as unequal crossing-over or gene conversion likely drive this homogenization, paralleling the well-characterized dynamics of rRNA arrays, histone gene clusters^25^, and certain highly repetitive neuropeptide loci (e.g., cnidarian neuropeptides^26^). All introns are phase 1 and display conserved lengths within species, with most introns being very short, measuring ~57 bp in *A. ceylanicum* and ~82 bp in both *A. caninum* and *A. duodenale* (Figure S4C), *underscoring the structural uniformity of these genes within species*.

By contrast, the chordate neurotensin/neuromedin N gene is encoded on four exons separated by introns of phase 1, 0, and 0, with the region encoding the neuromedin N and neurotensin peptides confined to a single exon. These observations, together with the distinct phylogenetic separation between chordates and nematodes, demonstrate that ancylotensins evolved convergently rather than through common ancestry or horizontal gene transfer from chordate neurotensin.

### Ancylotensins potently activate human and dog NTSR1

Most described biological effects of neurotensin are mediated through NTSR1^27^. NTSR1 is a versatile class A GPCR that preferentially couples to G_q/11_, leading to robust inositol phosphate (IP) signaling, but can also engage a wide range of Gα subtypes, including G_i/o_ and G_12/13_, and G_15_. To test whether ancylotensins could modulate mammalian NTSR1 signaling, we synthesized and tested one peptide from *A. caninum* (Aca-ATS1), one from *A. duodenale* (Adu-ATS1), and all three identified peptides from *A. ceylanicum* (Ace-ATS1-3). Several ancylotensins, including Aca-ATS1, contain a putative *O*-glycosylation site predicted using an O-glycan prediction tool (Figure S6). Accordingly, we also synthesized a glycosylated isoform, Aca-ATS1g (Ser9(Gal(β1-3)GalNAc)), for testing. In IPOne assays, all five ancylotensins displayed potent agonist activity at human NTSR1, with potencies and efficacies comparable to neurotensin (EC_50_ values: 0.53 nM (NTS), 0.24 nM (Aca-ATS1), 0.44 nM (Adu-ATS1), 0.16 nM (Ace-ATS1), 0.47 nM (Ace-ATS2), 1.18 nM (Ace-ATS3); Figure 3A, Table S2). The shorter peptide, Aca-ATS1s, which more closely resembles neuromedin N, and the glycosylated Aca-ATS1g both showed reduced activity compared to the non-glycosylated peptide (EC_50_ values: 1.75 nM (Aca-ATS1s) and 1.74 nM (Aca-ATS1g) Figure 3B and C, Table S2). All peptides also activated dog NTSR1 with similar potencies as observed at the human receptor when tested in the G_q_ protein dissociation assay (Figure 3D, Table S3). Aca-ATS1 and Aca-ATS1g were selected for more detailed pharmacological profiling at human NTSR1. In G protein dissociation assays, both isoforms coupled to multiple G protein subtypes and recruited β-arrestin 1 and 2, with the glycosylated peptide displaying overall lower activity while the non-glycosylated peptide shows more bias toward IP signaling over β-arrestin recruitment (Figure 3E-L, Figure S7, Table S4-7). Together, these findings demonstrate that ancylotensins mirror the pharmacological profile of host neurotensin, including its capacity to signal through diverse pathways.

**Figure 3.**
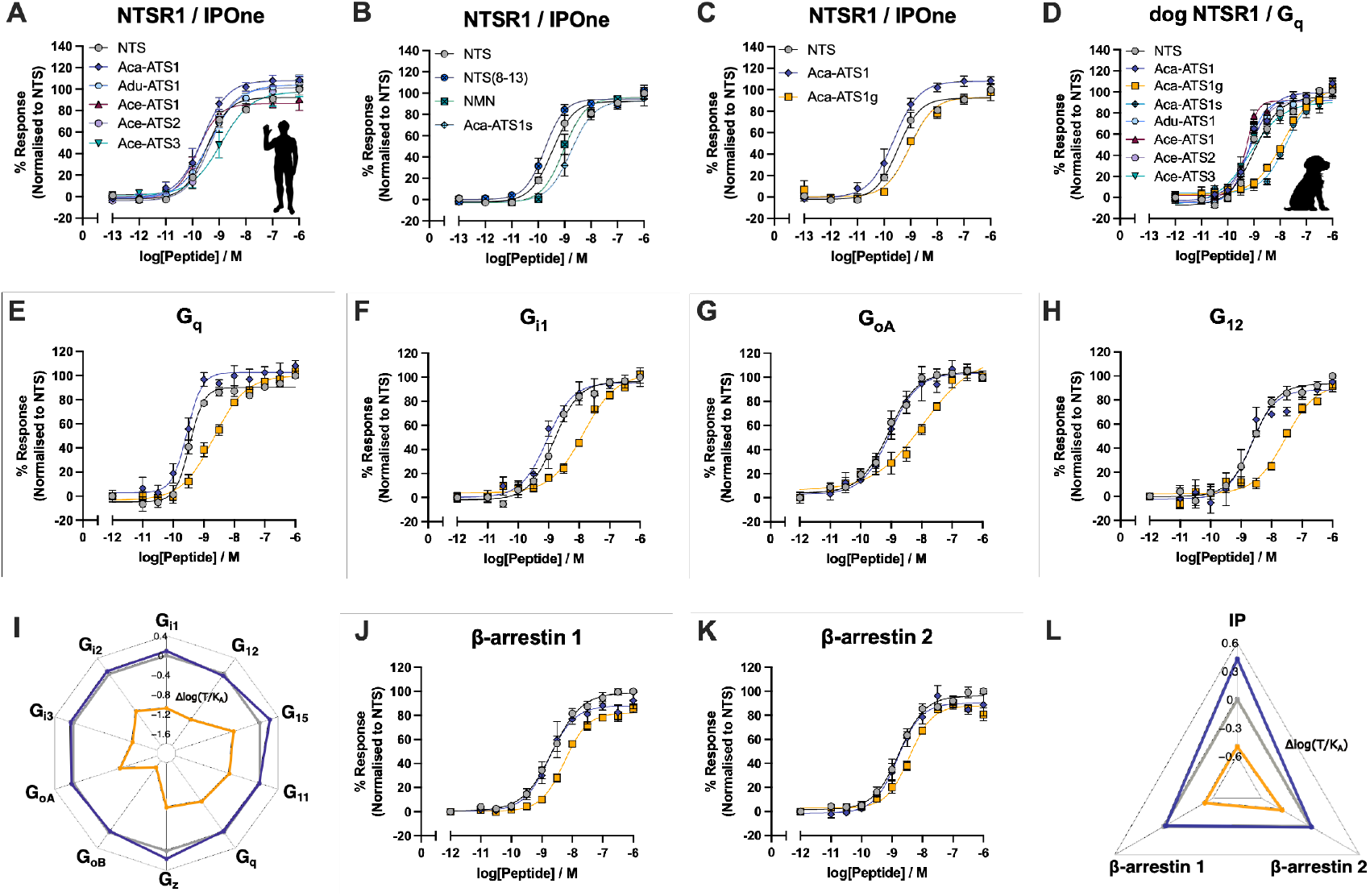
Ancylotensins potently activate human and dog NTSR1 and signal through various downstream pathways. (A-D) Representative dose-response curves of human NTS and NMN, and ancylotensins and ancylotensin isoforms (Aca-ATS1s and Aca-ATS1g) in mediating inositol monophosphate (IP) responses and G_q_ dissociation in HEK293T cells transiently transfected with human (A-C) and dog (D) NTSR1, respectively. (E-H) Representative dose-response curves of NTS, Aca-ATS1, and Aca-ATS1g in mediating dissociation of different G proteins. (I) G protein bias plot determined by Δlog(τ/K_A_) using NTS as the physiological reference ligand. (J-K) Representative dose-response curves of NTS, Aca-ATS1, and Aca-ATS1g in mediating β-arrestin 1/2 recruitment in HEK293T cells. (L) NTSR1 signaling bias plot determined by Δlog(τ/K_A_) using NTS as the physiological reference ligand. All error bars represent means ± S.E.M. of 3 biological replicates with technical duplicates.

### Ancylotensins mimic the core receptor binding motif of the mammalian hormone

Structural studies have defined how neurotensin and its shorter counterpart neuromedin N engage NTSR1^14,28^. Cryo-EM studies have demonstrated that ligand-bound NTSR1 can adopt two distinct active conformations, a canonical and a non-canonical state, likely reflecting flexibility in G protein coupling^14^. Binding occurs within the orthosteric pocket, with critical contacts formed by the conserved PYIL motif, particularly Tyr11 and Leu13, which interact with residues in transmembrane helices and extracellular loops. To determine whether ancylotensins structurally mimic these interactions, we resolved the high-resolution cryo-EM structure of the Aca-ATS1g-NTSR1 complex at 2.5 and 2.6 Å resolution (non-canonical and canonical orientation, respectively, accession numbers 9ZEA and 9ZE9, Table S8). We observed that Aca-ATS1g engages the receptor through the same critical contacts, particularly via the conserved PYIL motif and stabilizes both the canonical and non-canonical conformations (Figure 4A-C, Figures S8, S9, S10). Although the eight N-terminal residues of Aca-ATS1g are poorly ordered and show no clear contacts with the receptor, removing this region—as in the shorter analog Aca-ATS1s— substantially reduces NTSR1 activity (Figure 3B), suggesting that these residues form contacts that remain unresolved in our structure, likely due to their flexibility. Alanine mutations of the receptor core (*i*.*e*., at R6.55A, F6.58A, Y7.28A and W334 at the extracellular loop 3) reduce both activity of NTS and Aca-ATS1 and Aca-ATS1g by 100-1000-fold, in particular R6.54A, Y7.31A and Y7.35A are detrimental for NTS and ancylotensin signaling (Figure 4E, Table S9, Figure S11). These findings show that ancylotensins not only mimic host neurotensin pharmacologically but also reproduce its structural interactions at the atomic level.

**Figure 4.**
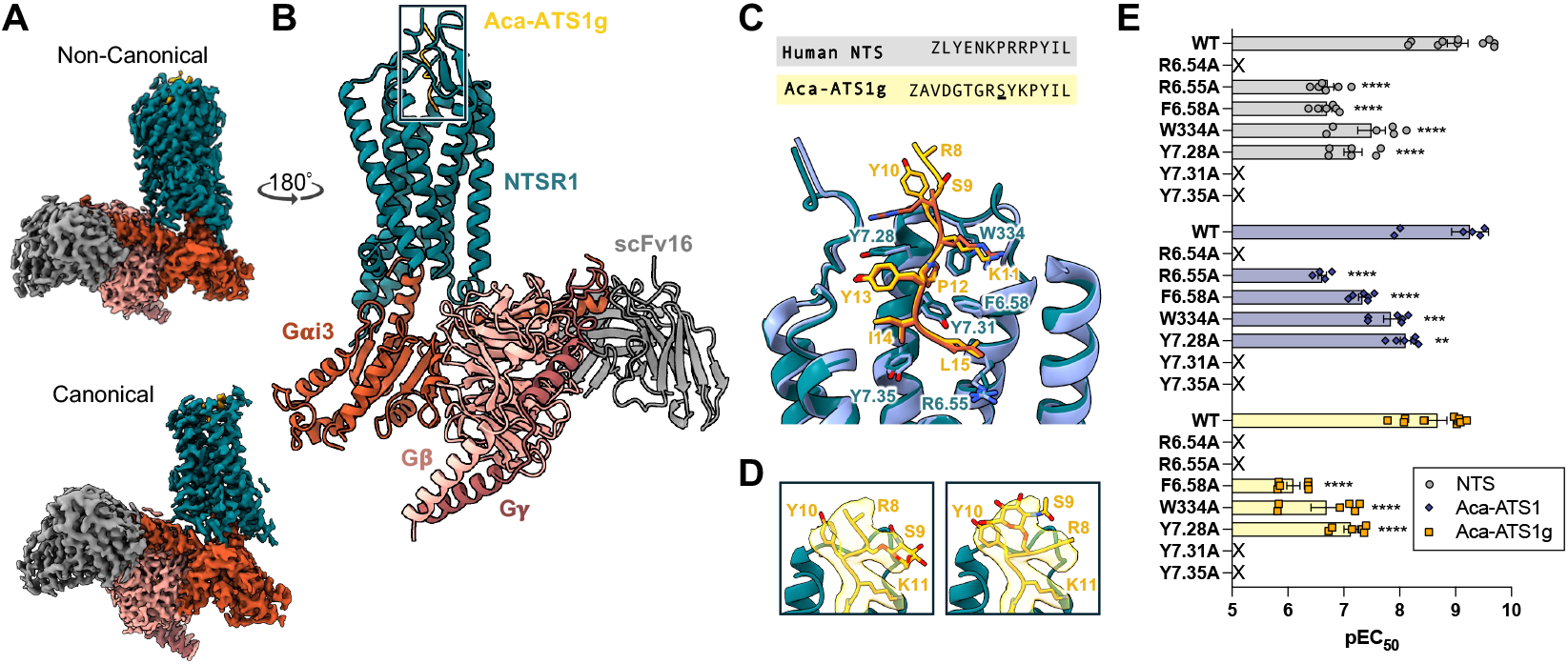
Ancylotensin Aca-ATS1g mimics the core receptor interactions of neurotensin at human NTSR1. (A) Cryo-EM maps of the Aca–ATS1g–NTSR1–Gi3 complex in non-canonical (top) and canonical conformations (bottom). (B) Cryo-EM structure of the non-canonical conformation. NTSR1 (dark teal), Aca-ATS1g (goldenrod), Gα_i3_ (burnt orange), Gβ (light pink), Gγ (dark rose) and scFv16 (gray). (C) Overlay of Aca-ATS1g–NTSR1 (gold and dark green) with human NTS(8-13)–NTSR1 (orange and blue; PDB: 4grv). Core Aca-ATS1g motifs (goldenrod) and receptor contact residues (dark teal, Ballesteros-Weinstein numbering^29^) are highlighted. Sequences of Aca-ATS1g and human NTS are shown above structure. Ser9 glycosylation in Aca-ATS1g underlined, Z represents pyroglutamate. (D) Unsharpened cryo-EM maps around the flexible Ser9 glycosylation site and Arg8 of Aca-ATS1g. (E) Scatter plot summarizing pEC_50_ values of how NTSR1 core mutations reduce NTS (grey), Aca-ATS1 (dark blue) and Aca-ATS1g (yellow) mediated IP accumulation. Error bars show mean ± S.E.M. of 3 biological replicates with technical duplicates. Statistical analysis was performed using One-way ANOVA, Dunnett test: **, p > 0.01, ***, p > 0.001, ****, p < 0.0001 and ns, non-statistically significant, vs. WT. X, non-determinable due to receptor inactivation.

### Ancylotensins modulate gut motility but are not analgesic

Administration of several neurotensin analogs, including a glycosylated neurotensin-like peptide from cone snail venom, contulakin-G, has been shown to provide central and peripheral analgesia in both dogs and humans^16,30,31^, with the glycosylation being important for the peptide’s *in vivo* efficacy^32^. Because hookworm larvae migrate through tissues before settling in the intestine, events that may be painful, and because adult hookworms attach to and feed on the intestinal mucosa, creating localized tissue damage, we hypothesized that ancylotensins may dampen nociceptive signaling. Such modulation could facilitate both silent tissue migration by larvae and blood feeding by adults. To test this, we examined the analgesic potential of Aca-ATS1 and Aca-ATS1g in mouse models of acute and post-surgical pain. Neither ancylotensin peptide exhibited detectable analgesic activity when administered at 10 mg/kg intraperitoneally (I.P.) in the hot plate assay (Figure 5A) or at 100 nmol intrathecally (I.T.) in the paw incision model (Figure 5B). For comparison, other neurotensin-like molecules, including contulakin-G and NT79, exert robust analgesic effects at much lower doses^31,33,34^.

**Figure 5.**
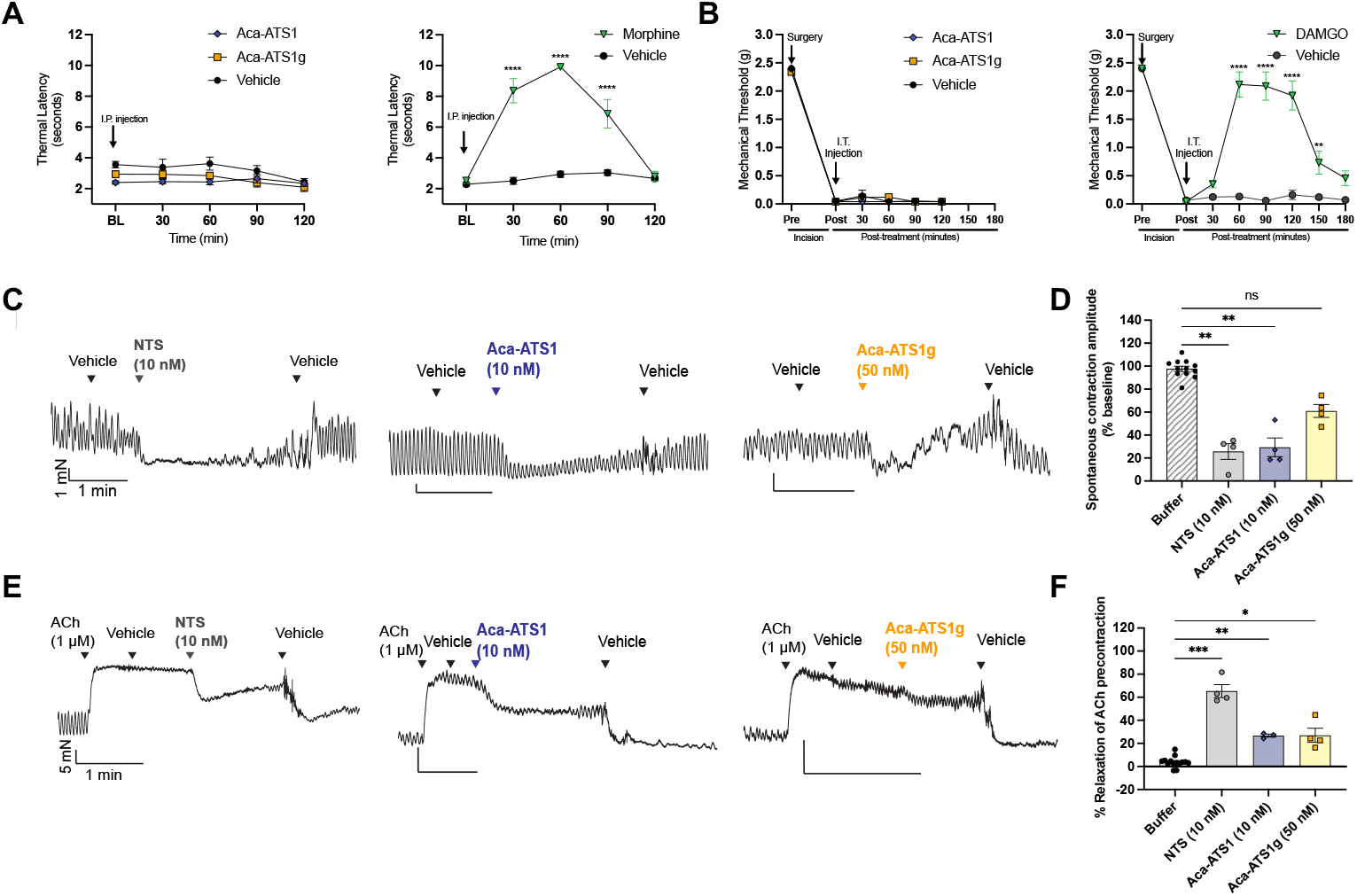
Ancylotensin Aca-ATS1 and Aca-ATS1g are not analgesic but modulate the contractility of the small intestine. (A) Analgesia was assessed in the hot plate assay in mice. Aca-ATS1, Aca-ATS1g (both at 10 mg/kg) and vehicle (saline) were given via intraperitoneal (I.P.) injection (left). Tail flick latency was measured for up to 120 minutes. At this concentration, neither peptide showed an effect when compared to vehicle. Morphine (10 mg/kg) was used as a positive control (right). (B) Central analgesia was assessed in the paw incision model with I.T. administration of 100 nmol Aca-ATS1 and Aca-ATS1g in saline. None of the peptides showed analgesic effects (left). DAMGO (0.1 nmol) was used as positive control (right). (C-D) NTS, Aca-ATS1 (10 nM) and Aca-ATS1g (50 nM) caused a reduction of the spontaneous contraction amplitude in *ex vivo* rat ileum. (C) Representative trace of each treatment. (D) Bar graphs of individual experiments. (E-F) NTS, Aca-ATS1 (10 nM) and Aca-ATS1g (50 nM) caused relaxation of acetylcholine (ACh)-induced ileum precontraction. (E) Representative trace of each treatment. (F) Bar graphs of individual experiments. Data represent technical replicates from 3-4 rats. Statistical analysis was performed using One-way ANOVA, Dunnett test. *, p < 0.05, **, p < 0.01, ***, p < 0.001, and ns, non-statistically significant.

Neurotensin is also known to influence intestinal smooth muscle tone in humans and dogs^35,36^. Because hookworms are blood-feeding parasites residing in the small intestine, modulation of gut contractility and local blood flow could directly aid in attachment and feeding. To test if ancylotensins can directly affect gut motility, we tested these peptides in *ex vivo* rat ileum preparations. NTS at 10 nM caused a marked reduction in spontaneous contraction amplitude to 25.7 ± 6.8% of baseline (*P* = 0.0034 versus buffer). Aca-ATS1 at 10 nM produced a similar effect, reducing amplitude to 29.4 ± 8.2% (*P* = 0.0021), while Aca-ATS1g at 50 nM reduced it to 70.0 ± 5.6% (*P* = 0.0021), with minimal effects at lower concentrations (Figure 5C-D). In acetylcholine (ACh)-precontracted tissue, NTS (10 nM) caused 65.4 ± 5.6% relaxation (*P* = 0.0004). Aca-ATS1 caused 26.8 ± 1.4% relaxation at 10 nM (*P* = 0.010) and 74.1 ± 10.8% at 50 nM (*P* = 0.036). Aca-ATS1g was less potent causing 27.0 ± 6.1% relaxation at 50 nM (*P* = 0.025) (Figure 5E-F). These findings indicate, like NTS, the hookworm-derived peptides can reduce intestinal contractility and promote smooth muscle relaxation, potentially creating a more favorable feeding environment.

## Discussion

Parasitic organisms evolve under strong selective pressures to establish, maintain, and exploit often long-term interactions with their hosts^37^. A central feature of this evolutionary process is the emergence of molecular strategies that evade detection and modulate host physiology in ways that favor parasite survival. One increasingly recognized strategy is the use of *doppelgänger peptides*—peptides that mimic the neuropeptides of another organism. Such molecular mimicry is well known in venomous animals and has also been documented in plant-parasite systems^38^, including root-knot nematodes that secrete phytohormone mimics to manipulate plant immunity and growth^39^, as well as in blood-feeding ectoparasites such as ticks^40^, mosquitoes^41^, and vampire bats^42^ that use host-hormone mimics to modulate inflammation and vascular function. However, whether endoparasitic animals similarly exploit neuropeptide mimicry has remained unexplored. Given that hookworms inhabit the mammalian intestine for years, we reasoned that prolonged residence in the gut might favor the evolution of hormone-like molecules that interfere with host gut–brain or enteric neuropeptide signaling systems to sustain a favorable physiological environment. Using computational discovery, we found that hookworms have convergently evolved neurotensin-like peptides, ancylotensins, that engage mammalian neurotensin receptors and modulate gut function.

It has recently been shown that prokaryotic gut residents, such as commensal bacteria, use molecular mimicry to shape host metabolic signaling. For example, microbial metabolites that mimic fatty-acid receptor ligands regulate metabolic hormones and glucose homeostasis^43^, and gut-bacteria-derived proteins that circulate in human blood improve metabolic markers in animal models^44^. Together, these lines of evidence suggest that endocrine mimicry may represent a common mechanism by which diverse gut inhabitants, from microbes to metazoan parasites, modulate host physiology to their advantage.

Our functional data demonstrate that ancylotensins do not provide analgesia but reduce intestinal contractility and promote smooth muscle relaxation, similar to the effects of host neurotensin. We propose that such modulation can facilitate both sustained blood feeding and stable intestinal occupancy by suppressing peristaltic movements, thereby reducing the likelihood of parasite expulsion and increasing local perfusion at the site of attachment. Given the pleiotropic functions of neurotensin, the role of ancylotensins may extend beyond gut contractibility. Hookworms rely on host-derived nutrients^1^, including blood lipids and glucose, and neurotensin is a key regulator of metabolic homeostasis. Parasitic worms are known to alter lipid and glucose metabolism^45^, and infection with *A. ceylanicum*, for example, induces hyperlipidemia in hamsters^46^. Although not examined here, ancylotensins could similarly influence neurotensin-linked metabolic^9,15^ or gut-brain pathways^8,11^.

While we have focused on doppelgänger peptides here, the repertoire of parasite-derived peptides that hijack host neuropeptide signaling systems is likely broader than currently recognized. Many such peptides may evade detection because they lack recognizable sequence motifs or are encoded by short open reading frames that fall below the thresholds of standard gene-prediction pipelines. Neuropeptide signaling pathways may be especially attractive targets for parasites because they are evolutionarily constrained. Host neuropeptides and their receptors share deep coevolutionary histories, making it difficult for hosts to evolve resistance without coordinated changes to both ligand and receptor. Doppelgänger peptides, including the ancylotensins described here, further limit escape by existing as multiple variants that occupy much of the receptor’s ligand-recognition space. To date, the only clear example of resistance to such mimicry is the decoy receptor XA21 in the rice species, *Oryza longistaminata*, which detects and neutralizes the bacterial PSY mimic RaxX50^47^. Independently of receptor-escape considerations, the presence of multiple ancylotensin copies, including identical variants, may reflect dosage effects: encoding repeated peptide sequences within a single precursor increases output from a single transcript, a strategy also seen in other peptide systems such as cnidarian neuropeptides^48^.

### Limitations of the study

Our study has several limitations. First, we did not confirm the presence of ancylotensins at the protein level, therefore the exact chemical nature of these peptides, including their processing sites and post-translational modifications, remains to be confirmed. Second, given the diverse physiological roles of neurotensin, ancylotensins may have additional functions that were not explored here. Finally, although we demonstrate that ancylotensins are potent agonists of dog and human NTSR1, we did not assess their activity at NTSR2, a closely related class A GPCR that is challenging to characterize *in vitro*, or at NTSR3 (also known as sortilin). Given the significant structural similarity between ancylotensins and neurotensin, it is likely that these peptides also interact with these receptors.

### Concluding remarks

Our findings reveal that hookworms have evolved neurotensin-like peptides that can modify host physiology, representing an example of molecular mimicry in a metazoan parasite. This strategy could provide multiple benefits: modulating local intestinal function to support feeding, while potentially influencing systemic metabolic circuits to optimize resource availability. More broadly, these findings highlight that parasite genomes may encode a rich but largely hidden repertoire of bioactive peptides that can interface directly with conserved host signaling pathways. As methods for detecting and characterizing small peptides improve, it is likely that many additional parasite peptides will be uncovered, offering new insights into host-parasite coevolution and potentially novel therapeutic targets.

## Acknowledgements

The support and resources from the Center for High Performance Computing at the University of Utah are gratefully acknowledged. We also thank Dr. Gaya P. Yadav for cryoEM data collection at the Laboratory for Biomolecular Structure and Dynamics (LBSD) of Texas A&M University. The LBSD is supported, in part, by the Department of Biochemistry & Biophysics, AgriLife, and the Texas A&M University. GPCRome screening of the kisspeptin-like peptide Cb-KP1 was generously provided by the National Institute of Mental Health’s Psychoactive Drug Screening Program, Contract # 75N95023C00021 (NIMH PDSP). This work was supported by a Villum Young Investigator Grant (19063 to H.S.-H.), and a Starting Grant from the European Commission (ERC-Stg 949830 to H.S.-H.), TLK is supported by an international postdoc fellowship from the Independent Research Fund Denmark (3102-00006B). KTS was supported by a Novo Nordisk Fond Hallas Møller Ascending Investigator grant (no. 0073793) and a Sapere Aude Research Leader Grant from the Independent Research Fund Denmark (2066-00043B). MJR is a CPRIT Scholar in Cancer Research supported by CPRIT award RR230042.

## Declaration of interest

The authors declare no competing interests.

## Methodology

### Database preparation

Expressed sequence tag (EST) data from *Ancylostoma* species were downloaded from NCBI and translated into putative open reading frames (ORFs) using EMBOSS getorf v6.6.0.0. These ORFs were combined with predicted proteomes of *A. caninum, A. duodenale*, and *A. ceylanicum, and N. americanus* obtained from WormBase ParaSite. To enrich for proteins unique to the parasitic Ancylostomidae lineage, we removed sequences with strong homology to *Caenorhabditis elegans* using MMseqs2 v18.8cc5c (E-value threshold 1e-4). We additionally assembled a dataset of proteins reported to be overexpressed during the parasitic L3 larval stage^19,20^. All sequences were assessed for secretory potential using a combination of SignalP 6^51^, DeepLoc^52^, and our custom non-classical secretion predictor SecEuk (described below). Secreted candidates were subsequently analyzed with DeepPeptide^23^ (with Esm2 model) to predict whether they encode short bioactive peptides.

### Eukaryotic non-classical secretion predictor, SecEuk

To construct a classifier for non-classically secreted proteins, we curated a positive dataset of 2,610 eukaryotic proteins annotated as secreted but lacking a signal peptide from SwissProt. As a negative dataset, we downloaded SwissProt-annotated non-secreted eukaryotic proteins (162,403 sequences) and subsampled these to match the positive dataset’s size and length distribution. The combined dataset was partitioned into training, validation and test sets (70/20/10%) using GraphPart^53^, ensuring <40% pairwise sequence identity across partitions.

We implemented SecEuk using the esm2_t6_8M_UR50 protein language model with a sequence-classification head in the HuggingFace EsmForSequenceClassification framework. Models were trained using the training parameters below: fp16=True, evaluation_strategy=“epoch”, learning_rate=5e-5, per_device_train_batch_size=4, per_device_eval_batch_size=4, num_train_epochs= 4, weight_decay= 0.99, max_grad_norm=0.5, save_total_limit=2, metric_for_best_model=“accuracy”, and gradient_accumulation_steps=2. During training, ESM-2 embeddings were frozen for the first four epochs and then unfrozen for a final fine-tuning epoch. Training used the AdamW optimizer with automatic mixed precision (FP16) to accelerate training and reduce memory usage. Model selection and evaluation were performed on the held-out validation and test sets; the final SecEuk model achieved 0.72 accuracy **and** 0.81 ROC–AUC on the independent test set and showed improved ROC–AUC over DeepLoc^**52**^ on the validation partition. However, both SecEuk and DeepLoc predictions were retained and integrated for downstream secretion calls to increase confidence.

For reproducibility, all training scripts, model checkpoints, hyperparameter files and evaluation logs are available on HuggingFace (see Data & Code Availability). Training was performed on GPU hardware.

### Doppelgänger toxin identification

To identify Ancylostomidae peptides mimicking vertebrate neuropeptides, we curated a reference panel of 53 human neuropeptides and retrieved homologs from other vertebrates using BLASTP searches against the SwissProt database. Mature peptide regions and adjacent propeptide processing sites were extracted, aligned with MAFFT v7.526, and profile hidden Markov models (HMMs) were constructed using HMMER hmmbuild. These HMMs were searched against the assembled Ancylostomidae sequence database, and significant matches were manually evaluated for domain structure, signal peptide presence, and conservation of key processing motifs.

### Molecular evolutionary analyses

We identified homologs of the initial ancylotensin hit using tblastn and blastp querying the ancylostome genome assemblies and predicted protein sequences, respectively. The gene structures of the identified genes were determined with exonerate using both protein to genome (p2g) and nucleotide to genome (e2g) models. The exon sequences identified with exonerate were extracted and aligned using MAFFT^56^, and a maximum likelihood exon tree was reconstructed using IQ-TREE^58^ v3.0.1 with seed 52392 on a single thread. The K2P model of evolution was chosen with ModelFinder based on the Bayesian information criterion, and bootstraps were calculated with UFboot with 1000 replicates.

The gene neighborhood twenty genes up- and down-stream of the ancylotensin genes were retrieved from the NCBI genome annotation files, and orthology between the genes were determined with CLANS^63^.

Macrosyntenic relationships were determined by first scaffolding the *A. caninum and A. ceylanicum* genomes using RagTag^60^ with *A. duodenale* as the model. Homologous genes were identified with OrthoFinder^62^, and these were visualized using the R library macrosyntR^61^.

### Glycosylation prediction

We predicted potential *O*-glycosylation sites of Ser/Thr residues in the ancylotensin-encoding precursors using the online version of NetOGlyc v4.0 (https://services.healthtech.dtu.dk/services/NetOGlyc-4.0/)^57^.

### Peptide synthesis and purification

Peptides were either purchased from commercial vendors or synthesized in-house (to >90 % purity) as described below. Aca-ATS1 and Aca-ATS1s were synthesized by solid-phase peptide synthesis in 0.1 or 0.05 mmol scale using preloaded Fmoc-protected Tentagel R HMPA resin from Rapp Polymere (Tuebingen, Germany) for C-terminal carboxylic acid sequences and TentaGel® S RAM Resins for C-terminal amide sequences. Fmoc-protected amino acids, coupling reagents, and solvents used for the synthesis were purchased from Iris Biotech (Marktredwitz, Germany). The synthesis was performed on a Syro I instrument from Biotage (Uppsala, Sweden). Coupling conditions were room temperature (RT) for 2 × 120 min using 5.2 equiv amino acids, 4.7 equiv N-[(1H-benzotriazol-1-yl)(dimethylamino) methylene]-N-methylmethanaminium hexafluorophosphate N-oxide, 5.2 equiv 1-hydroxy-7-azabenzotriazol, and 8 equiv N,N-diisopropylethylamine in dimethylformamide (DMF) relative to resin. Deprotection was performed at RT with 40% piperidine in DMF for 3 min followed by 20% piperidine in DMF for 15 min. Washing steps were with 2 × N-methyl-2-pyrrolidone, 1 × dichloromethane (DCM), and 1 × DMF. After completion of the peptide assembly, the peptidyl resin was washed with 3 × DCM and dried. The peptide was released from approximately 0.025 mmol resin using a mixture (2 mL) of 95% trifluoroacetic acid (TFA), 2.5% triethylsilyl, and 2.5% water over 2.5 h. Cold diethylether (13 mL at −20 °C) was added, and the mixture was further cooled to −85 °C for 30 min before the peptide was isolated by centrifugation. The peptide was redissolved in a mixture of TFA, water, and acetonitrile (ACN) and purified using 1 out of 2 methods: Either the purification happened on a Dionex Ultimate 3000 HPLC system (Thermo Fisher, Waltham, USA) using a Luna C18(2) column from Phenomenex (Torrance, USA, 5 μm, 100 Å, 250 × 10 mm). Alternatively, the purification happened on a Selekt System (Biotage, Sweden) using a 50 g Biotage Sfär Bio C18 D column. For both systems, a gradient of 5–100% ACN in water/0.1% formic acid was applied. The product was isolated and lyophilized by freeze-drying. The final product was analyzed by liquid chromatography (LC)–mass spectrometry (MS) on a Dionex Ultimate 3000 ultrahigh-performance LC system from Thermo Fisher connected to an Impact HD mass spectrometer (Bruker, Bremen, Germany). Adu-ATS1, Ace-ATS2, and Ace-ATS3 were synthesized on a Gyros PurePep® Chorus peptide synthesizer on a 50 µmol scale. The linear peptides were constructed on preloaded Fmoc-Leu-Wang resin (Advanced ChemTech, cat. no. SL5145) (0.36 mmol/g loading). Deprotection of Fmoc was achieved with 20% piperidine in DMF for 2 min twice. Amide coupling was achieved by using 6 equivalents of amino acid (200 mM in DMF), 6 equivalents of HATU (200 mM in DMF) and 12 equivalents of NMM (600 mM in DMF) for 25 min. Cleavage of the linear peptides was achieved by treatment of the peptidyl-resin with TFA/H_2_O/TIPS (TFA: trifluoracetic acid, TIPS: triisopropylsilane) (95:2.5:2.5) (8). Side-chain protecting groups for amino acids were as follows: Lys and Trp, tert-butyloxycarbonyl (Boc); Ser, Thr, and Tyr, tert-butyl ether (tBu); Cys and Asn, trityl (Trt); Arg, 2,2,4,6,7-pentamethyl-dihydrobenzofuran-5-sulfonyl (Pbf); Glu, tert-butyl ester (OtBu); Asp, 5-butylnon-5-yl ester (OBno). All crude peptides were purified with a 0.1% TFA buffered H_2_O (buffer A) and 0.1% TFA buffered 9:1 ACN:H_2_O (buffer B) gradient on an Agilent 1260 Infinity II HPLC system using either a Phenomenex Jupiter C12, 4 µm (250 x 21.2 mm) (prod. numb. 00G-4396-P0-AX) preparative column or Phenomenex Jupiter C4, 10 µm (250 x 21.2 mm) (prod. numb. 00G-4168-P0-AX) preparative column at 15 mL.min^-1^. Fractions collected from HPLC were analyzed by LC-MS on a Phenomenex Jupiter C12, 4 μm (50 × 2 mm) (prod. numb. 00B-4396-B0) column at 0.5 mL.min^-1^ with a 0.1% formic acid buffered H_2_O (buffer A) and 0.1% formic acid buffered 9:1 ACN:H_2_O (buffer B) gradient on an Agilent 1260 Infinity II Quadrupole LC-MS system. Fractions containing targeted product (based on LC-MS) were collected and lyophilized. Purity of isolated peptides were determined by peak integration of blank subtracted HPLC spectra. Ace-ATS1, the kisspeptin-like peptide from *Caenorhabiditis bovis* (Cb-1) was synthesized commercially by Genscript. The glycosylated isoform Aca-ATS1g was synthesized commercially by Polypeptide (USA) using glycosylated Fmoc-L-Ser amino acids synthesized by CarboSyn (USA, Fmoc-L-Ser (Gal β(1-3)GalNAc)-OH, peracetate. All peptides were validated by HPLC and ESI-MS prior to use.

### Chemical reagents

Unless otherwise specified, all chemical and cell culture reagents were sourced from Sigma and ThermoFisher respectively. Neurotensin, neurotensin(8-13), neuromedin N and Ace-ATS1 were custom-synthesized by Genscript. Larger amounts of the non-glycosylated (Aca-ATS1) and glycosylated Aca-ATS1g were further custom-synthesized by GenScript and Polypeptide (San Diego, USA), respectively. Fmoc-L-Ser(Gal(β1-3)GalNAc) was purchased from CarboSyn (USA). The mass and purity (> 95 %) of the peptides were verified by RP-HPLC and MS prior to use.

### Cell culture

HEK293T cells were given by Prof. Hans Braüner-Osborne (University of Copenhagen, Denmark) and were cultured in DMEM supplemented with 10% FBS (Gibco) and 100 U/mL penicillin and 100 µg/ml streptomycin at 37°C in a humidified incubator supplemented with 5 % CO_2_. Mycoplasma testing was conducted regularly to ensure the lack of mycoplasma contamination.

### DNA constructs

3xHA-tagged human NTSR1 construct was obtained from cDNA.org (cDNA resource center, Bloomsburg PA, US). Constructs for N-terminus tagged with 3xHA Dog-ntsr1 (*Canis lupus familiaris*) (sequence accession number: XM_038434300.1) inserted in pcDNA3.1(+) plasmid were custom-synthesized by Twist Biosciences. cAMP sensor using YFP-Epac-RLuc (CAMYEL)^49^ was given kindly by Prof. Hans Braüner-Osborne (University of Copenhagen, Denmark). mVenus-β-arrestin 1/2 constructs were kind gifts from Dr. Tao Che (Washington University, St. Louis). C-terminally NanoLuciferase (NLuc)-tagged human NTSR1 inserted in pcDNA3.1(+) used in the β-arrestin recruitment studies were customed synthesized by GenScript. TRUPATH G protein constructs were gifts from Prof. Bryan Roth (University of North Carolina, Chapel Hill) (Addgene kit #1000000163). All NTSR1 mutants (i.e. R6.54A, R6.55A, F6.58A, W334A, Y7.28A, Y7.31A and Y7.35A) inserted in pcDNA3.1(+)-N-terminal-HA used in this study were custom synthesized by GenScript. All DNA constructs were sequenced and confirmed prior to use.

### IPOne Gq assay

HEK293T cells were seeded onto 6-well plates at a density of 900,000 cells/well. Cells were transfected the next day with 3xHA-tagged human NTSR1, dog-ntsr1 or NTSR1 mutants (all at 1 µg) using Polyfect (Qiagen) according to manufacturer’s protocol. 48 h post-transfection, HEK293T cells were split using 0.05% Trypsin-EDTA solution and cell density was determined using an automated cell counter. Manufacturer’s protocol were then largely followed in next steps. Cells were resuspended in stimulation buffer that contains lithium chloride provided by the IPOne Gq activation kit (Revvity). Cells were seeded at 40,000 cells / well on white 384 Optiplate (Revvity) and were stimulated with peptide ligands for 1 hour at room temperature. Equal volume of Eu- and Tb-tagged antibodies diluted in lysis buffer provided by assay kit were added on the cells and were incubated for 1 h. HTRF signal was determined using PHERAstarFSX plate reader (BMG Labtech). HTRF ratio was determined by dividing 665 nm signal over 620nm signal.

### β-arrestin 1/2 recruitment assay DNA constructs

HEK293T cells were seeded onto 6-well plates at a density of 900,000 cells/well. Cells were transfected the next day with C-terminally-NLuc-tagged human NTSR1 (0.3 µg) and mVenus-β-arrestin 1/2 constructs (0.3 µg) at 1:1 ratio using Polyfect (Qiagen) according to manufacturer’s protocol. 16 to 24 h post-transfection, HEK293T cells were seeded onto white 96 well plate (Grenier Bio-one) coated with PLL at a density of 40,000 cells/well and were allowed to grow overnight. On the day of assay, cells were washed once with PBS and 80 µL of assay buffer which composed of PBS with 0.5 mM MgCl_2_, adjusted to pH 7.4, were added onto each well. 10 µL of NLuc substrate furimazine (Promega) were added onto each well and incubated for 3 min. 10 µL of 10x concentrated ligands were then added manually onto the plate and BRET signal was measured instantaneously using PHERAstarFSX plate reader (BMG Labtech) equipped with BRET1 475 (30 nm bandwidth) and 535 (30 nm bandwidth) filters continuously for 30 min.

### TRUPATH G protein dissociation assay

The following protocol was adopted from^64^. In brief, HEK293T cells were seeded onto 6-well plates at a density of 900,000 cells/well. Cells were transfected the next day with 3xHA-tagged human NTSR1 or NTSR1 mutants (0.3 µg), RLuc8-tagged G protein constructs (0.2 µg), pcDNA3.1-Gβ3 (0.2 µg), pcDNA3.1-Gγ2-GFP2 (0.2 µg) (at a ratio of 1.5:1:1:1 and the total DNA amount per well was 0.9 µg) using Polyfect (Qiagen) according to manufacturer’s protocol. 16 to 24 h post-transfection, HEK293T cells were seeded onto white 96 well plate (Grenier Bio-One) coated with poly-L-lysine (PLL) at a density of 40,000 cells/well and were allowed to grow overnight. On the day of assay, cells were washed once with PBS and 80 µL of assay buffer which composed of PBS with 0.5 mM MgCl_2_, adjusted to pH 7.4, were added onto each well. 10 µL of Rluc8 luciferase substrate Prolume Purple (Nanolight coorperate) were added onto each well and incubated for 5 min. 10 µL of 10x concentrated ligands were then added manually onto the plate and BRET signal was measured instantaneously using PHERAstarFSX platereader (BMG Labtech) equipped with BRET2 410 (80 nm bandwidth) and 515 (30 nm bandwidth) filters continuously for 30 min.

### CAMYEL biosensor assay

HEK293T cells were seeded onto 6-well plates at a density of 900,000 cells/well. Cells were transfected the next day with 3xHA-tagged VPAC1R, VPAC2R or PAC1nR (0.15 µg) and CAMYEL biosensor^49^ (0.30 µg) (at a ratio of 1:2 the total DNA amount per well was 0.45 µg) using Polyfect (Qiagen) according to manufacturer’s protocol. 16 to 24 h post-transfection, HEK293T cells were seeded onto white 96 well plate (Grenier) coated with poly-L-lysine (PLL) at a density of 40,000 cells/well and were allowed to grow overnight. On the day of assay, cells were washed once with phosphate-based saline (PBS) and 80 µL of assay buffer which composed of PBS with 0.5 mM MgCl_2_, pH 7.4, were added onto each well. 10 µL of Coelentrazine-h luciferase substrate (Nanolight) were added onto each well and incubated for 5 min. 10 µL of 10 x ligands were then added manually onto the plate and BRET signal was measured instantaneously using PHERAstarFSX platereader (BMG Labtech) equipped with BRET1 475 (30 nm bandwidth) and 535 (30 nm bandwidth) filters continuously for 30 min.

### BRET ratio calculation

BRET1 ratio was determined by dividing the BRET signal at 535 nm over that at 475 nm. BRET2 ratio was determined by dividing the BRET signal at 515 nm over that at 410 nm. The net BRET ratio was determined by subtracting the BRET ratio with drugs with the BRET ratio of vehicle control. Net BRET ratio at 30-min time points for each ligand concentration tested were determined and plotted against log(concentration) of drugs tested. Results of test ligands were normalized as percentage of maximal neurotensin response. Concentration response curves were fitted with a 4-parameter logistic equation (equation 1) using GraphPad Prism (v.10.6.0):

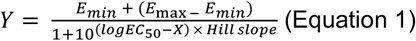

### G protein bias quantification

The methods used in^65^ to quantify G protein bias was largely followed. In brief, neurotensin was chosen to be the physiological reference ligand given its physiological significance as well as its full agonism at human NTSR1. The operational model (equation 2) was used to determine the transduction ratios (τ/K_A_).

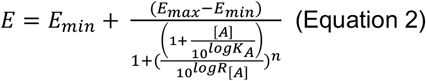

where E is the effect of the ligand, [A] is the concentration of test ligands, E_max_ is the maximal response of the system, E_min_ is the minimum response of the system, logK_A_ is the logarithm of the equilibrium dissociation constant, n is the slope of the transducer function, log R is the logarithm of the transduction ratio (τ/K_A_). The operational model was manually input in GraphPad Prism (v.10.6.0) according to ^65^. Each experiment was analyzed using the operational model which all individual experiments were fitted with a global shared slope. The LogR (equivalent to log[τ/K_A_]) values were estimated by GraphPad Prism (v 10.6.0) and were used to calculate the relative efficacy (RE) using equation 3 and 4.

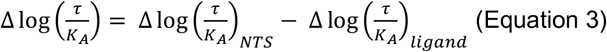

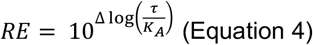

S.E.M. of 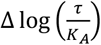, were calculated using the following equation 5:

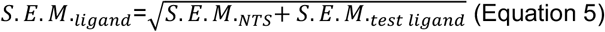

### Expression and purification of NTSR1

Pellets containing NTSR1 were expressed in Sf9 cells and snap-frozen in liquid nitrogen. To start purification of the receptor, the cell pellets were thawed and lysed in a hypotonic lysis buffer containing 10 mM HEPES at pH 7.5, 1 mM EDTA, 1 mM benzamidine, Pierce Universal Nuclease, 1 mM MgCl_2_, protease inhibitor cocktail, and 100 μM TCEP, and stirred at 100 rpm for 1 hour at 4°C. Subsequently, the membranes were harvested via ultracentrifugation at 100,000xg for 35 minutes. The pellets were resuspended in a solubilization buffer containing 20 mM HEPES pH 7.5, 500 mM NaCl, 1 mM MgCl_2_, 1 mM benzamidine, Pierce Universal Nuclease, protease inhibitor cocktail, and 100 μM TCEP. Detergent was added dropwise to the resuspended pellets to a final concentration of 1% LMNG/0.1% CHS/0.1% Sodium Cholate, while stirring at 4°C. After gently stirring for 3 hours at 4°C, ultracentrifugation was done at 100,000xg for 35 minutes to remove insoluble debris. The supernatant was pooled and supplemented with 20 mM imidazole, then loaded over a TALON resin column that had been washed with 10 column volumes of a buffer containing 20 mM HEPES pH 7.5, 20 mM imidazole, 500 mM NaCl, and 0.1% LMNG/0.01% CHS. The protein was eluted by loading 2 column volumes of buffer containing 20 mM HEPES pH 7.5, 250 mM imidazole, 250 mM NaCl, 0.01% LMNG/0.001% CHS, and 10% glycerol over the column. Purified NTSR1 was concentrated down to 500 μL, then subjected to SEC chromatography with a Superdex 200 column in buffer containing 20 mM HEPES pH 7.5, 5% glycerol, 100 mM NaCl, and 0.01% LMNG/0.001% CHS. Peak fractions were collected and pooled together, concentrated to a lower volume, supplemented with 5% glycerol, and snap-frozen with liquid nitrogen for storage and use with complexation.

### Expression and purification of dominant-negative G_**i3**_

Purification of dominant negative (DN) G_i3_ heterotrimer expressed in *T*.*ni*. cells began with thawing and resuspension in lysis buffer containing 20 mM HEPES pH 7.5, 1 mM MgCl_2_, 5 mM β-mercaptoethanol, 1 mM EDTA, 5% glycerol, 100 µM GDP, protease inhibitor cocktail, and Pierce Universal Nuclease, and stirred for 30 minutes at 4°C. The resulting lysate was harvested via ultracentrifugation at 100,000xg for 35 minutes. Pellets were resuspended in a solubilization buffer containing 20 mM HEPES pH 7.5, 1 mM MgCl_2_, 5% glycerol, 100 mM NaCl, 1% sodium cholate, Pierce Universal Nuclease, 100 µM GDP, protease inhibitor cocktail, and 5 mM β-mercaptoethanol. After gently stirring at 4°C for 1 hour, ultracentrifugation was done for 35 minutes at 100,000xg, and the supernatant was pooled and supplemented with 30 mM imidazole and incubated with Ni-NTA beads for 1 hour on ice. Nickel beads were harvested via centrifugation at 300 xg and packed into a chromatography column. The column was washed with a series of washes of 10 column volumes with buffers containing 30 mM imidazole with 50% solubilization/50% E2 buffer, 25% solubilization/75% E2 buffer, 12.5% solubilization/87.5% E2 buffer, and 100% E2 buffer (E2 buffer contained 20 mM HEPES pH 7.5, 100 µM GDP, 1 mM MgCl_2_, 5% glycerol, 100 mM NaCl, 5 mM β-mercaptoethanol, and 0.05% LMNG/0.005% CHS). To elute DN G_i3_ from the column, 3 column volumes of E2 buffer and 250 mM imidazole were loaded over the column, followed by incubation in a 3,500 MWCO dialysis bag with 1 mg of 3C protease per 50 mg of G-protein, for overnight dialysis at 4°C. The next day, the Ni-NTA column was washed with 10 column volumes of E2 buffer and 30 mM imidazole. The overnight cleavage was then loaded over the column, and flow through was collected. Two additional column volumes of E2 buffer and 30 mM imidazole were washed over the column, and flow through was collected. The resulting G-protein was concentrated down to 500 µL in a 30 kDa MWCO spin concentrator, and was subjected to SEC chromatography using a Superdex 200 column in a buffer with 20 mM HEPES pH 7.5, 20 µM GDP, 100 µM TCEP, 100 mM NaCl, 1 mM MgCl_2_, 5% glycerol, and 0.01% LMNG/0.001% CHS.

### Expression and Purification of scFv16

scFv16 was purified as described previously^66^ and purified scFv16 was concentrated to 13 mg/mL and snap frozen with liquid nitrogen in buffer containing 20 mM HEPES pH 7.5, 100 mM NaCl, and 15% glycerol for future use.

### Formation and purification of NTSR1-Aca-ATS1g-G-protein-scFv16 complex

NTSR1 and Aca-ATS1g were incubated together on ice for 1 hour. To begin complex formation, the Aca-ATS1g-bound receptor was mixed with a molar excess of G-protein and incubated for 1 hour on ice. Apyrase was added to remove GDP and HRV 3C protease was added to cut the GFP tag off of NTSR1, followed by overnight incubation on ice. Complex was diluted with buffer of 5-fold more volume consisting of 20 mM HEPES pH 7.5, 100 μM Aca-ATS1g, 100 mM NaCl, and 0.01% LMNG/0.001% CHS. The diluted complex was loaded over an M1 flag resin column, then washed with buffer containing 20 mM HEPES pH 7.5, 10 μM Aca-ATS1g, 100 mM NaCl, and 0.005% LMNG/0.0005% CHS. NTSR1-Aca-ATS1g-G-protein complex was eluted from the column with a buffer of 20 mM HEPES pH 7.5, 10 μM Aca-ATS1g, 1 mM EDTA, 100 mM NaCl, flag peptide, and 0.005% LMNG/0.0005% CHS. The eluent was concentrated to a lower volume and loaded onto a size exclusion Superdex 200 column with a buffer of 20 mM HEPES pH 7.5, 10 μM Aca-ATS1g, 0.001% LMNG/0.0001% CHS/0.00033% GDN, 100 mM NaCl, and 2 mM MgCl_2_. Peak fractions of the complex were pooled and concentrated to ~10 mg/ml for cryo-EM analysis.

### Cryo-EM sample prep and imaging

Cryo-EM samples were prepared with a Vitrobot Mark IV held at 100% humidity and 4°C by application of 3 μL of sample to UltrAuFoil R1.2/1.3 300 mesh grids, glow discharged in a Pelco unit for 45 seconds at 10 mA, and plunge frozen in liquid ethane. Grids were imaged on a G4 Titan Krios at 300 kV with a K3 direct electron detector and a Bioquantum energy filter. Details of the imaging parameters can be found in Table S8.

### Data processing and model building

All data were processed in cryoSPARC^67^ beginning with patch motion correction and patch CTF estimation. Template-based picking was performed with existing GPCR-G-protein complex templates, followed by particle extraction and 2D classification to remove junk particles. Additional particle cleaning was done with rounds of *ab-initio* model generation with multiple classes and heterogeneous refinement. Final particles were reconstructed with initial non-uniform refinement, CTF refinement and reference-based motion correction followed by final non-uniform and receptor/Ras domain local refinement. Initial models for the receptor were derived from PDB:6OSA and 6OS9^14^ while dominant negative Gi3 was obtained from PDB:7T10^68^. Manual model building was performed in Coot^69^ with iterative rounds of real space refinement in Phenix^70^. Details of the imaging, final maps and models can be found in Table S8.

### Intestinal contractility assay

Male Wistar rats (250-300 g) were killed by CO_2_ asphyxiation and cervical dislocation. Segments of ileum (10–20 mm) were attached to wire tissue holders, mounted in isolated organ baths under 5-10 mN resting tension and maintained at 37ºC in Tyrode’s solution. Isometric contractions were measured using an ADI MLT0420 0-20 g force transducer connected to an ADI FE221 bridge amp and 26 series Powerlab and recorded using LabChart v8.1.21 (ADInstruments). Preparations were equilibrated for at least 30 min before the addition of drugs. Experiments involving the use of rat tissue were approved by the University of Queensland Animal Ethics Committee (tissue sharing from animals used in pharmacology teaching lab) (UQ AEC; approval number 2023/AE000409) and conducted in accordance with the Australian Code of Practice for the Care and Use of Animals for Scientific Purposes, 8th edition (2013).

### Mice behavioral experiments

Before any behavioral experiment or testing, the animals were brought to the testing room in their home cages for at least 1 hour for acclimation. Testing always occurred within the same approximate time of day between experiments, and environmental factors (noise, personnel, and scents) were minimized. All testing apparatus (cylinder, grid boxes, etc.) were cleaned between uses. The experimenter was blinded to treatment group by another laboratory member delivering coded drug vials, which were then decoded after collection of all data

### Tail flick thermal latency

Pre-injection tail-flick baselines were determined in a 52°C water bath using male and female CD-1 (ICR) mice (5 - 10 weeks) with 10 second cutoff time. Injections of Aca-ATS1 and Aca-ATS1g at 30 mg/kg and vehicle (sterile water) were given via intraperitoneal (I.P.) injection and tail flick latencies were determined over a time course; 30-, 60-, 90-, and 120 min post injection. A 150-min time point was taken if the animal did not return to baseline 120 min post injection. One animal was excluded from the study as it had an abdominal injury from injection. Two-way ANOVA and Tukey’s multiple comparison with the vehicle control were performed.

### Paw incision and mechanical allodynia assessment

Mechanical thresholds were determined before surgery using calibrated Von Frey filaments (Stoelting Co.) with the “up-down” method and four measurements after the first response per mouse. The size range of stimuli was between 2.44 (0.4 mN) and 4.56 (39.2 mN). The starting filament was 3.61 (3.9 mN). The filament was placed perpendicular to the skin with a slowly increasing force until it bent; it remained bent for approximately 1 s and was then removed. Data were analyzed using the nonparametric method of Dixon, as described by Chaplan et al.^71^. The mice were housed in a Von Frey apparatus (Bioseb invivo instruments) with Plexiglas walls and ceiling and a wire mesh floor. The surgery was performed by anesthesia with ~2-5% isoflurane in 2% oxygen, preparation of the right plantar hind paw, sterilized with iodine and 70% ethanol 3 times, and a 5-mm incision made through the skin and fascia with a no. 10 scalpel. The muscle was elevated with curved forceps, leaving the origin and insertion intact, and the muscle was split lengthwise using the scalpel to not sever the tendon. 300 μL 1 mg/kg gentamycin was then administered. The wound was then closed with 5-0 nylon sutures. The next day, the mechanical threshold was again determined as described above, and Aca-ATS1, Aca-ATS1g, and vehicle (sterile water) were given via intrathecal (I.T.) injection at 100 nmol dose (in a volume of 5 μL). No animals were excluded from these studies. Mechanical threshold compared to vehicle at both the 30-, 60-, 90-, and 120-min time points were measured. A Two-way ANOVA with multiple comparisons was performed. Results correspond to means ± SEM.

### Quantification and Statistical Analyses

All statistical analyses, fittings, and plotting of curves were performed using GraphPad Prism 10. For GPCR assays, experiments were conducted in triplicate, and EC_50_ and EC_50_ ± S.E.M. were calculated in Prism. For the paw incision model, the effects at different time points were analyzed using two-way ANOVA with Geisser-Greenhouse correction and Dunnett’s post-test for multiple comparisons (all compared to vehicle), while the area under the curve plot was analyzed using one-way ANOVA with Dunnett’s post-test for multiple comparisons (all compared to vehicle). For the SNI model, the effect over time was analyzed as described for the paw incision model (all compared to the vehicle on the ipsilateral side), while the area under the curve was analyzed using a two-way ANOVA with Dunnett’s post-test for multiple comparisons (all compared to the vehicle on the ipsilateral side).

### Images and Visual Representation

All graphs were created using GraphPad Prism 10. All images that are not data graphs or plots were created using Adobe Illustrator, and Inkscape. Fig. 1A was illustrated by ilusea studio.

## Notes

### Competing Interest Statement

The authors have declared no competing interest.

