## Supporting Figures for "Mimics of the chordate gut–brain hormone neurotensin in parasitic intestinal hookworms"

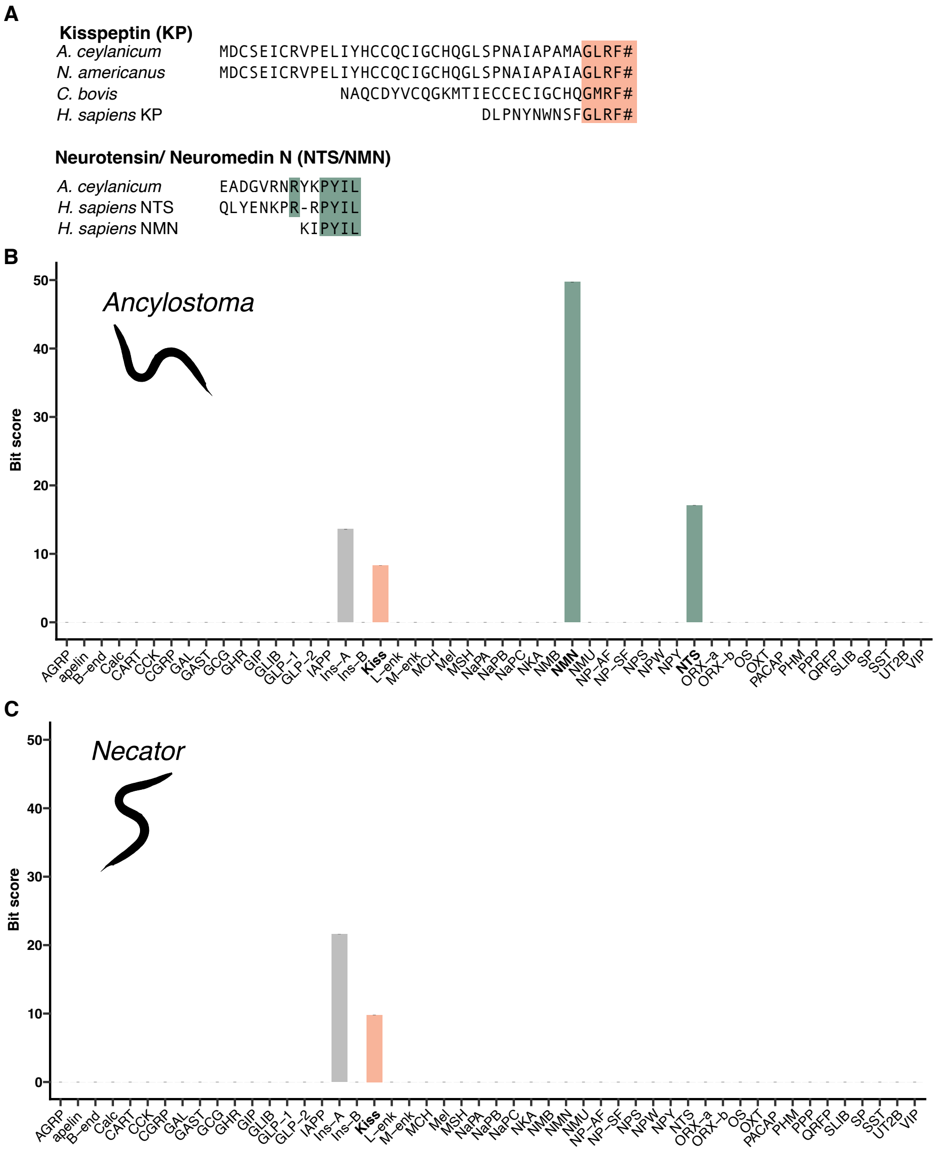


**Figure S1**. Identification of pHMM hits in the hookworm secreted protein database. (A) Sequence alignments of hookworm doppelgänger peptide candidates and their corresponding human hormone counterparts. Identical amino acids are highlighted; # denotes C-terminal amidation. An additional kisspeptin-like peptide was identified in *Caenorhabditis bovis* (Cb-KP1) by homology. (B) Hidden Markov model (HMM) searches in the Ancylostoma database recovered hits corresponding to kisspeptin (orange), neurotensin/neuromedin N (dark green), and the insulin A-chain. Further investigation showed that the insulin A-chain hits have homologs in other, non-parasitic nematodes and were therefore excluded, as they are not unique to hookworms. (C) HMM searches in *Necator americanus* similarly recovered a kisspeptin-like peptide and insulin A-chain hits, the latter of which were excluded for the same reasons as in Ancylostoma.

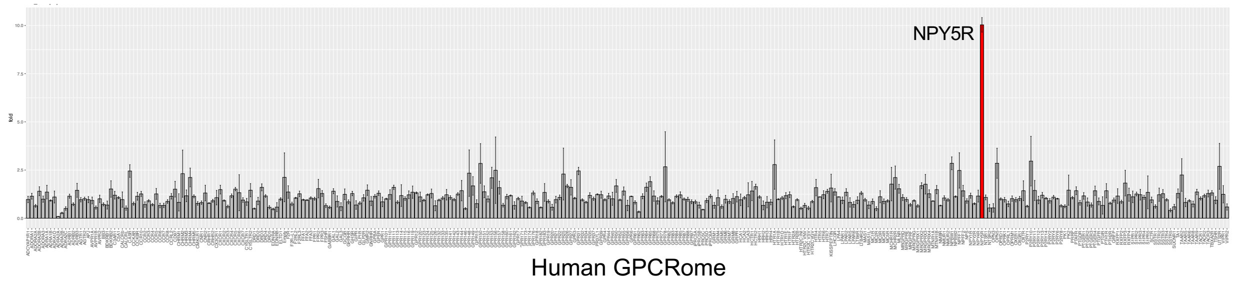


**Figure S2.** GPCRome screen of kisspeptin-like peptide. Attempts to synthesize the kisspeptin-like sequence from *Ancylostoma ceylanicum* failed and the other kisspeptin-like peptide, Cb-KP1, from *Caenorhabiditis bovis* was custom synthesized by Genscript instead. The synthetic peptide was sent to the National Institute of Mental Health's Psychoactive Drug Screening Program (PDSP) at the University of North Carolina at Chapel Hill for G protein-coupled receptor-ome (GPCRome) screening using the PRESTO-Tango assay. At 10 μM Cbo-KP1 showed a 10-fold increase in basal levels (~4 relative luminescence units) at the neuropeptide Y5 receptor (NPY5R). Dopamine receptor DRD2 with 100 nM quinpirole was used as an assay control. X axis depicts the 318 human GPCRs tested in this assay. When tested in-house using the same PRESTO-Tango assay, Cb-KP1 was found to be a weak, partial agonist of the NPY5R, not providing a full dose-response curve when tested at up to 300 μM (data not shown).

**Neurotensin/neuromedin N (*Homo sapiens*)**
MMAGMKIQLVCMLLLAFSSWSLCSDSEEEMKALEADFLTNMHTSKISKAHVPSWKMTLLNVCSLVNNLNSPAEETGEVHEEELVA
RRKLPTALDGFSLEAMLTIYQLHKICHSRAFQHWELIQEDILDTGNDKNGKEEVIKRKIPYILKRQLYENKPRRPYILKRDSYYY

**Neurotensin/neuromedin N (*Canis lupus familiaris*)**
*MMAGMKIQLVCMILLAFSSWSLC*SDSEEEMKALEADLLTNMHTSKISKASVSSWKMTLLNVCSFVNNLNSQAEETGEFREEELITR
RKFPTALDGFSLEAMLTIYQLQKICHSRAFQQWELIQEDVLDAGNDKNEKEEVIKRKIPYILKRQLYENKPRRPYILKRGSYYY

**Full length ancylotensin precursor sequences**
***Ancylostoma caninum***
MWASIALLLFANVVCSFERQAAEEPRRSWKPYILRQVAEEAGRSWKPYILKRSECGHVALAVLPFSVAV
***Ancylostoma caninum***
MWASIALLLFANVVCSFERQAAEEPRRSWKPYILRQAAEERRRSWKPYILREADEETRKSREPYILRQVAEEAGRSWKPYILRQV
AEEAGRSWKPYILRQVAEEAGRSWKPYILRQVAEEAGRSWKPYILRQVAEEAGRSWKPYILRQVAEEAGRSWKPYILRQVAEEA
GRSWKPYILRQAAEETSRNWKPYILSMFSQLFILIHFASFLEHLA
***Ancylostoma ceylanicum***
MLLILYIFFLHASGSDEREVDGHGGPLQILVSKREADGVRNRYKPYILKREADGVRNRYKPYILKREADGVRNRYKPYILKREADGV
RNRYKPYILKREADGVRNRYKPYILKREADGVRNRYKPYILKREADGVRNRYKPYILKREADGMLV
***Ancylostoma duodenale***
MWATIALLLFANVVCSFGREADVQPRNPYKPYILKREADEENRNRWKPYILKREADEETRNRWKPYILKREADEVNRNRWKPYIL
KREADEETRNRWKPYILKREADEETRNRWKPYILKRETDEENRNRWKPYILKRETDEENRNRWKPYILKREADEENRNRWKPYIL
KREADEENRNRWKPYILKREADEENRNRWKPYILKREADEENRNRWKPYILSMFNHLFILIHFASFLEHLSL

**Partial ancylotensin precursor sequences**
***Ancylostoma caninum***
ERQAADGTGRSYKPYILKRQAVDGTGRSYKPYILKRQAADGTGRSYKPYILKRQAVDGTGRSYKPYILKRQAVDGTGRSHKPYIL
KRQAVDGTGRSYKPYILKRQAVDGTGRSYKPYILSKCGHVAFAVLPFSATV
***Ancylostoma duodenale***
EREADGTPRNPYKPYILKREADGTPRNPYKPYILKREADGTPRNPYKPYIL
***Ancylostoma ceylanicum***
GLHDFGTRRSWKPYILKRQVAEEAGRSWKPYILKRQVAEEAGRSWKPYILKRQAAEETGRNWKPYILKRQVAEEAGRSWKPYIL
KRQVAEEAGRSWKPYILKRQVAEEAGRSWKPYILKRQVAEEAGRSWKPYILKRQVAEEAGRSWKPYILKRQVAEEAGRSWKPYIL
KRQVAEEAGRSWKPYILKRQVAEEAGRSWKPYILKRQVAEEAG
***Ancylostoma ceylanicum***
SEREADGVRNRYKPYILKREADGVRNRYKPYILKREADGVRNRYKPYILKREADGVRNRYKPYILKREADGVRNRYKPYILKREAD
GVRNRYKPYILKREADGVRNRYKPYILS

**Figure S3.** Ancylotensin precursor sequences identified in this study in comparison with human and dog neurotensin/neuromedin N. Signal sequences are shown in pink, proteolytic cleavage sites in red, neuromedin N in yellow, and neurotensin and ancylotensin in orange.

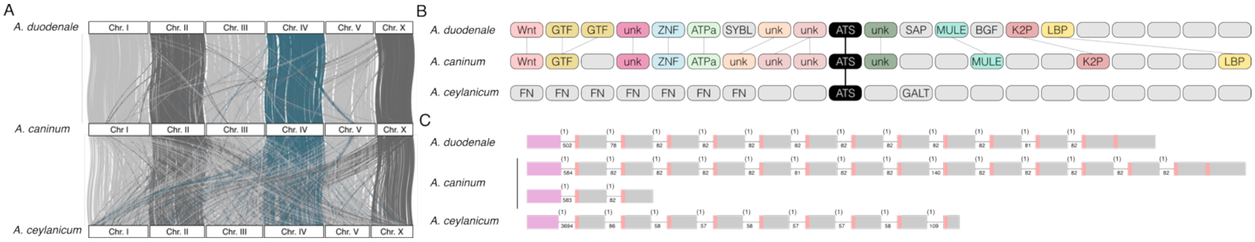


**Figure S4.** Synteny analysis and gene structures of ancylotensins from *A. duodenale*, *A. ceylanicum*, and *A. caninum*. (A) Macrosynteny analyses between the genomes of the three ancylostome species show a large degree of homology in chromosome structure among the three species. The ancylostensin genes are located within chromosome IV highlighted in dark green. (B) Genomic neighborhood of ancylotensin genes on chromosome IV show conserved synteny between *A. duodenale* and *A. ceylanicum*. Each box represents a protein encoding gene. Genes with significant homology between the three species are colored and when possible annotated. Uncolored and empty boxes represents proteins of unknown function with no homology within the genomic loci (C) Gene structure of identified full-length ancylotensin genes with exons represented by boxes and introns by lines. Signal sequence shown in pink, and predicted cleavage sites in red. Introns are all phase 1 (shown as (1)), and have conserved lengths, shown below the lines in base pairs.

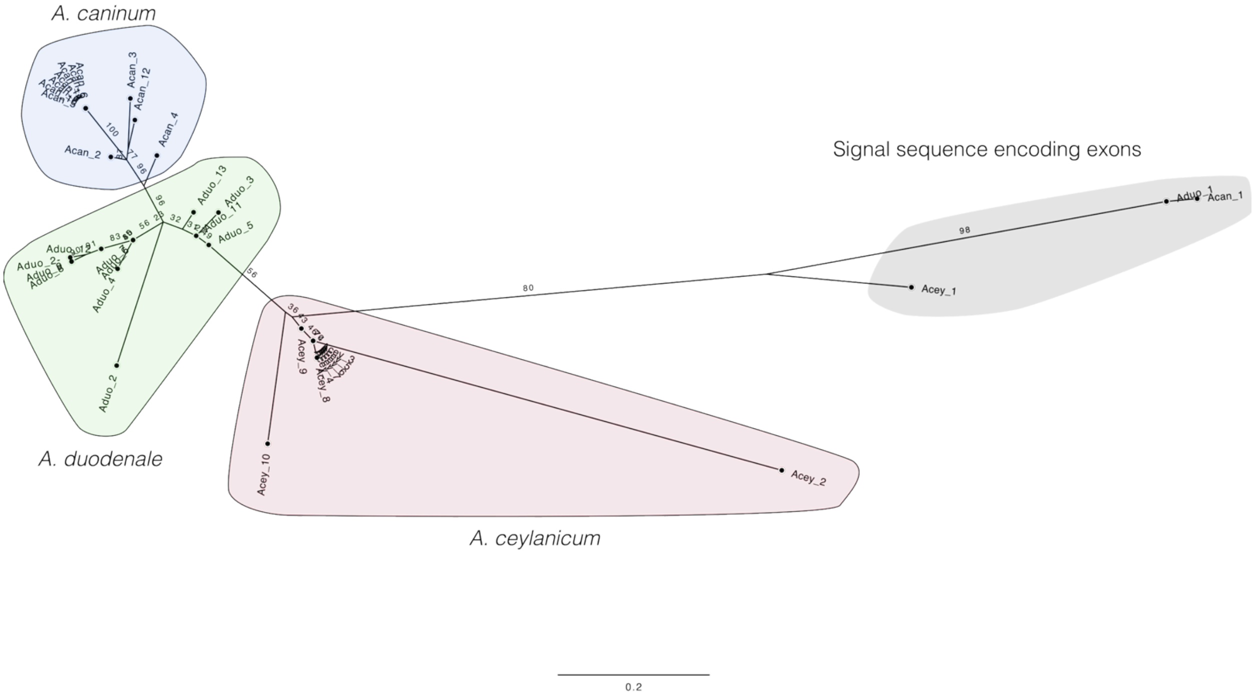


**Figure S5.** Ancylotensin repeat phylogeny show higher similiarity of the exons within species than between. Maximum-likelihood phylogeny (K2P model based on Bayesian Information Criterion) of ancylotensin-encoding exons reveals strong evidence of exon homogenization within species: exons cluster primarily by species rather than by orthology, indicating that repeated units within each genome are more similar to one another than to their counterparts in other species.

*A. duodenale* MWATIALLLFANVVCSFGREADVQPRNPYKPYILKREADEENRNRWKPYILKREADEETRNRWKPYILKREADEVNRNRWKPYILKR
EADEETRNRWKPYILKREADEETRNRWKPYILKRETDEENRNRWKPYILKRETDEENRNRWKPYILKREADEENRNRWKPYILKR
EADEENRNRWKPYILKREADEENRNRWKPYILKREADEENRNRWKPYILSMFNHLFILIHFASFLEHLSL

EREADGTPRNPYKPYILKREADGTPRNPYKPYILKREADGTPRNPYKPYIL

*A. caninum* MWISIALLLFANVVCSFEREADEETRKSREPYILKRQVAEEAGRSWKPYILKRQVAEEAGRSWKPYILKRQVAEEAGRSWKPYILKR
QVAEEAGRSWKPYILKRQVAEEAGRSWKPYILKRQVAEEAGRSWKPYILKRQVAEEAGRSWKPYILKRQVAEEAGRSWKPYILKR
QVAEEAGRSWKPYILKRQVAEEAGRSWKPYILKRQVAEEAGRSWKPYILKRQVAEEAGRSWKPYILKRQVAEEAGRSWKPYILKR
QAAEETSRNWKPYILSMFSQLFILIHFASFLEHLAR

QAADGTGRSYKPYILKRQAVDGTGRSYKPYILKRQAADGTGRSYKPYILKRQAVDGTGRSYKPYILKRQAVDGTGRSHKPYILKR
QAVDGTGRSYKPYILKRQAVDGTGRSYKPYILSKCGHVAFAVLPFSATV

MWASIALLLFANVVCSFERQAAEEPRRSWKPYILRQVAEEAGRSWKPYILKRSECGHVALAVLPFSVAV

*A. ceylanicum* MLLILYIFFLHASGSDEREVDGHGGPLQILVSKREADGVRNRYKPYILKREADGVRNRYKPYILKREADGVRNRYKPYILKR
EADGVRNRYKPYILKREADGVRNRYKPYILKREADGVRNRYKPYILKREADGVRNRYKPYILKREADGMLV

**Figure S6.** Predicted glycosylation sites within ancylotensin precursors suggests that some ancylotensins are glycosylated. Predicted *O*-glycosylation sites (serines) are show are shown as blue residues as determined by NetOGlyc-4.0 with prediction values above 50 %. Predicted peptides are shown in orange. The signal peptides are shown in green.

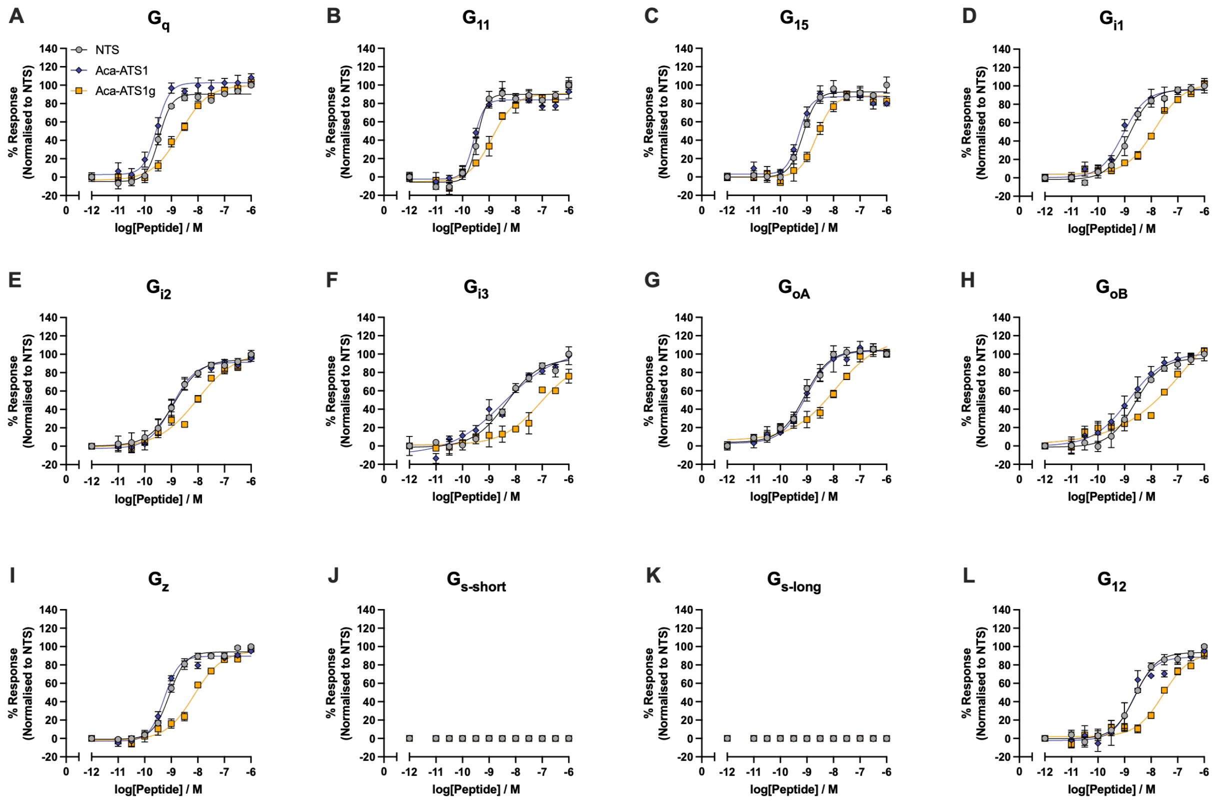


**Figure S7.** Representative dose-response curves of NTS, Aca-ATS1, and Aca-ATS1g in mediating dissociation of different G proteins in HEK293T cells transiently transfected with human NTSR1. Error bars represent means ± S.E.M. of 3 biological replicates with technical duplicates.

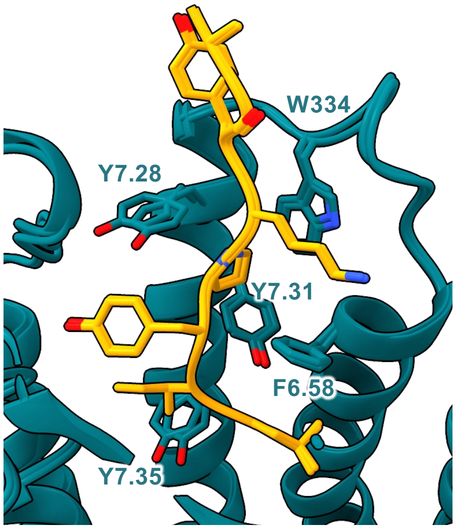


**Figure S8.** Ancylotensin-binding interface for the canonical and non-canonical conformation structures. Overlay of the structures of the core ancylotensin residues for the non-canonical and canonical conformation structures with Aca-ATS1g (goldenrod), NTSR1 (dark teal).

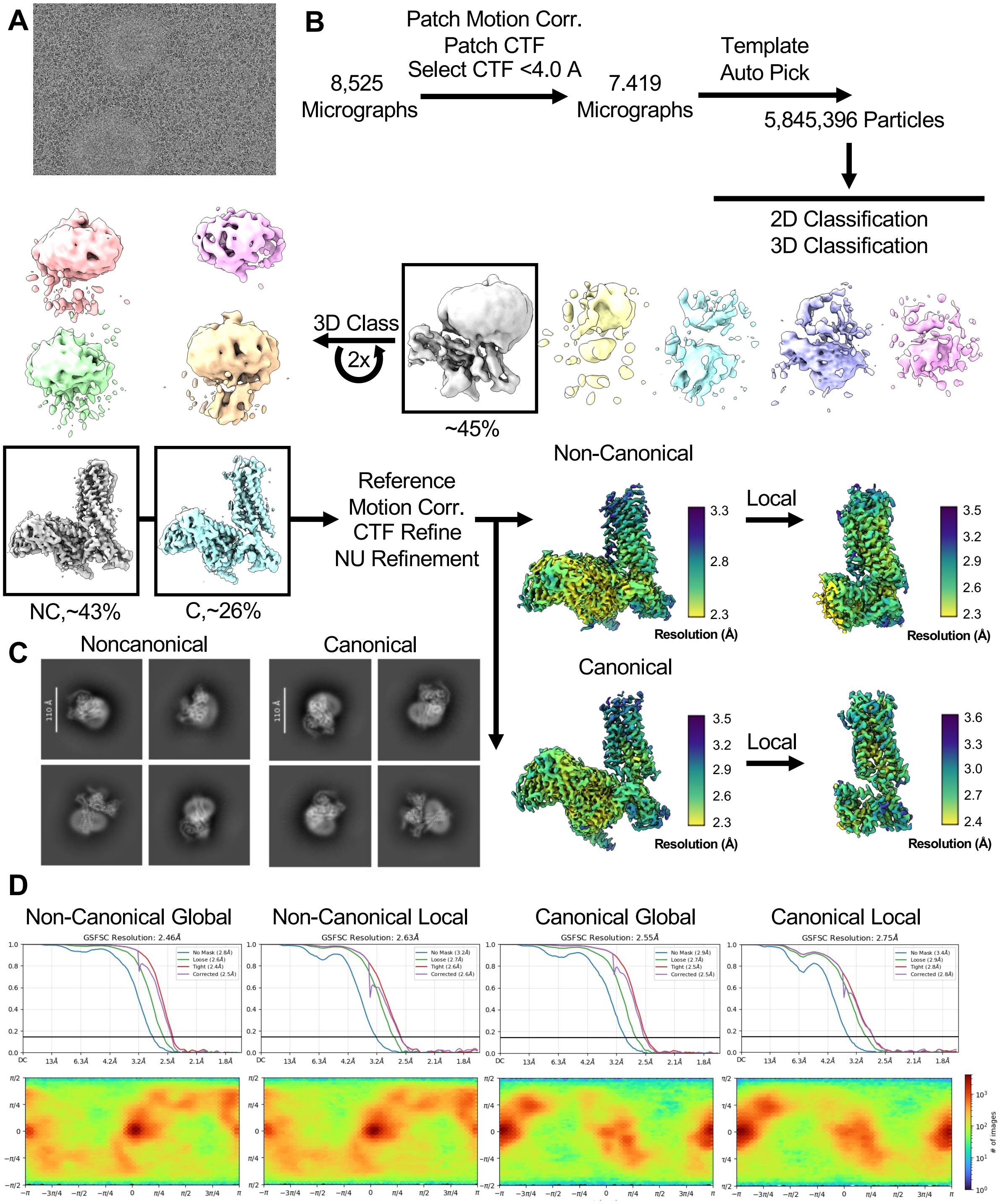


**Figure S9.** Cryo-EM data processing. **(**A) Representative cryo-EM micrograph of NTSR1 complex (B) Cryo-EM processing workflow employed in this study including final maps with local resolution. C) 2D Class averages of the particles contributing to the two final structures in this work. D) FSC plots and Euler angle distributions for the maps derived in this work.

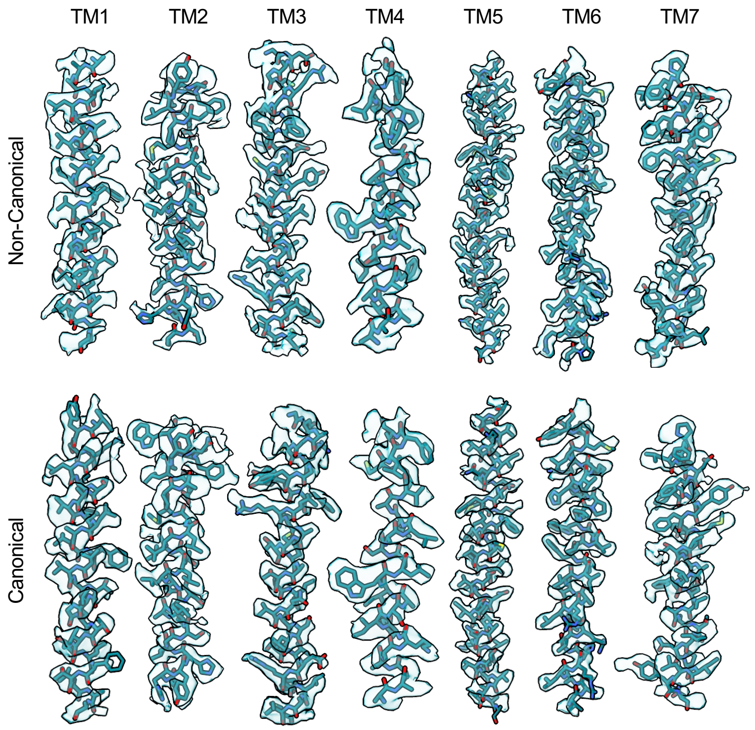


**Figure S10.** Map-Model agreement and the local refined receptor maps. Map-Model agreement for the NTSR1 transmembrane helices and the local refined receptor maps for the canonical and non-canonical states resolved in this work.

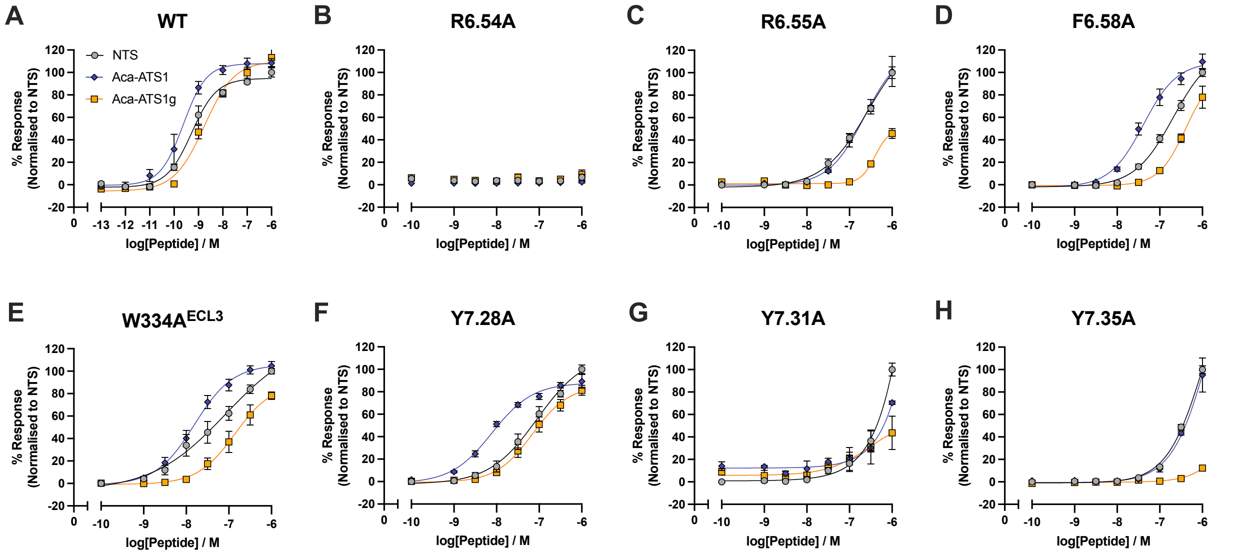


**Figure S11.** Core receptor mutation data. Dose response curves showing how NTSR1 core mutations (naming according to Ballesteros-Weinstein numbering) reduce IP accumulation mediated by NTS, Aca-ATS1 and Aca-ATS1g. Error bars represent means ± S.E.M. of 3 biological replicates with technical duplicates.
